# Deciphering Transcription in *Cryptosporidium parvum*: Polycistronic Gene Expression and Chromatin Accessibility

**DOI:** 10.1101/2025.01.17.633476

**Authors:** Rui Xiao, Rodrigo P. Baptista, Fiifi Agyabeng-Dadzie, Yiran Li, Yinxin Dong, Robert J. Schmitz, Travis C. Glenn, Jessica C. Kissinger

## Abstract

Once considered rare in eukaryotes, polycistronic mRNA expression has been identified in kinetoplastids and, more recently, green algae, red algae, and certain fungi. This study provides comprehensive evidence supporting the existence of polycistronic mRNA expression in the apicomplexan parasite *Cryptosporidium parvum*. Leveraging long-read RNA-seq data from different parasite strains and using multiple long-read technologies, we demonstrate the existence of defined polycistronic transcripts containing 2-4 protein encoding genes, several validated with RT-PCR. Some polycistrons exhibit differential expression profiles, usually involving the generation of internal monocistronic transcripts at different times during development. ATAC-seq in sporozoites reveals that polycistronic transcripts usually have a single open chromatin peak at their 5-prime ends, which contains a single E2F binding site motif. Polycistronic genes do not appear enriched for either male or female exclusive genes. This study elucidates a potentially complex layer of gene regulation with distinct chromatin accessibility akin to monocistronic transcripts. This is the first report of polycistronic transcription in an apicomplexan and expands our understanding of gene expression strategies in this medically important organism.

**Significance:** *Cryptosporidium parvum* is a parasite that causes human gastrointestinal disease, profoundly impacting infants and immunocompromised individuals worldwide. *C. parvum* isolation, culturing, cloning, and its compact 9.2 Mb genome pose challenges for research. We have discovered that, unlike most eukaryotes, which express one gene per mRNA, *C. parvum* exhibits widespread polycistronic transcription, whereby 10% of its protein encoding genes are expressed as multiple genes per single mRNA molecule. Mining the published *C. parvum* single-cell atlas with proxy sequences for polycistronic expression, we observed patterns of polycistron expression that vary with developmental stages, suggesting a role in the parasite’s lifecycle. By illuminating this unique aspect of *C. parvum* biology, our findings provide novel insights into *Cryptosporidium* gene expression.

## Introduction

*Cryptosporidium parvum*, an early-branching member of the phylum Apicomplexa(1, 2), is a medically important eukaryotic pathogen impacting human health globally(3-5). It is challenging to treat as there is only one approved drug that it not recommended for the most vulnerable, and there is no vaccine. Cryptosporidiosis poses a severe threat, particularly to infants(6) and the immunocompromised, where it can lead to life-threatening watery diarrhea(5). In healthy individuals, the infection can still lead to severe, but self-limiting diarrhea and cramping for two to six weeks(4). The limited treatment options(3, 4) for infants and the immunocompromised underscore an urgent need for novel approaches to understand and combat this parasite(4).

Despite recent advances in *Cryptosporidium* genetics(7-11) and *in-vitro* culturing(12-14), *Cryptosporidium*’s regulatory network underpinning gene expression remains poorly characterized. *Cryptosporidium*’s genomic landscape is characterized by its remarkable compactness(10, 11), consisting of just 9.259 million base pairs and housing 3,923 protein-encoding genes along with 913 small and long non-coding RNA genes(11, 15-17). The transcriptional landscape, inferred from strand-specific short-read RNA-seq evidence, shows extensive untranslated region (UTR) overlap between genes, regardless of their coding strand. The organism’s inability to *de novo* synthesize purines and pyrimidines(18, 19) and its complete reliance on nucleotide salvage pathways may be a contributor to this compactness. Our exploration of this intricate transcriptional terrain employed single-molecule sequencing to refine the transcript boundaries within a newly assembled telomere-to-telomere (T2T) genome sequence of *C. parvum* IOWA(11). This approach uncovered 201 polycistrons (PTs), which contain 2-4 protein-encoding genes each. The 201 PTs account for approximately 10% of the organism’s protein-encoding potential. This finding of polycistronic transcripts in *C. parvum* is unprecedented within the Apicomplexa and sheds light on a previously unrecognized layer of gene regulatory complexity.

Polycistronic transcription is a well-documented phenomenon in prokaryotes that is usually known as operons, where multiple genes’ coding sequences (CDS) are transcribed into a single mRNA transcript(20). It has also been documented in certain eukaryotic lineages such as *C. elegans*, and the dinoflagellates but it is most extensive in the kinetoplastids(21-23). Eukaryotic polycistronic transcription typically involves a *trans*-splicing mechanisms, where individual open reading frames (ORFs) have a spliced-leader sequence added onto the 5′ end of the processed transcript before the ORF can be translated(24). However, the type of polycistronic transcription we observed in *C. parvum* does not align with *trans*-splicing paradigms, rather, it shows similarities to the non-*trans*-splicing polycistronic mechanisms reported in some fungi(25, 26), *Drosophila*(27), and, more recently, in green(28) and red algae(29). These polycistronic transcripts, which are also often accompanied by the co-expression of internal monocistrons (IMCs) (Fig.1A), represent a putative functional unit of unknown significance to the parasite’s gene regulatory and developmental processes. This study delves into the transcriptional and regulatory nuances of the polycistronic transcripts in *C. parvum*, employing an integrative approach that combines single-molecule long-read RNA-seq for validation, global bulk ATAC-seq for chromatin accessibility, short-read RNA-seq for developmental stage-based expression profiling, and gene ontology enrichment analysis for functional insights. Our findings illuminate the complex landscape of transcriptional regulation in *C. parvum*. Our comprehensive analysis suggests that *C. parvum* likely possesses a non-*trans*-splicing polycistronic gene expression system.

**Fig 1.**
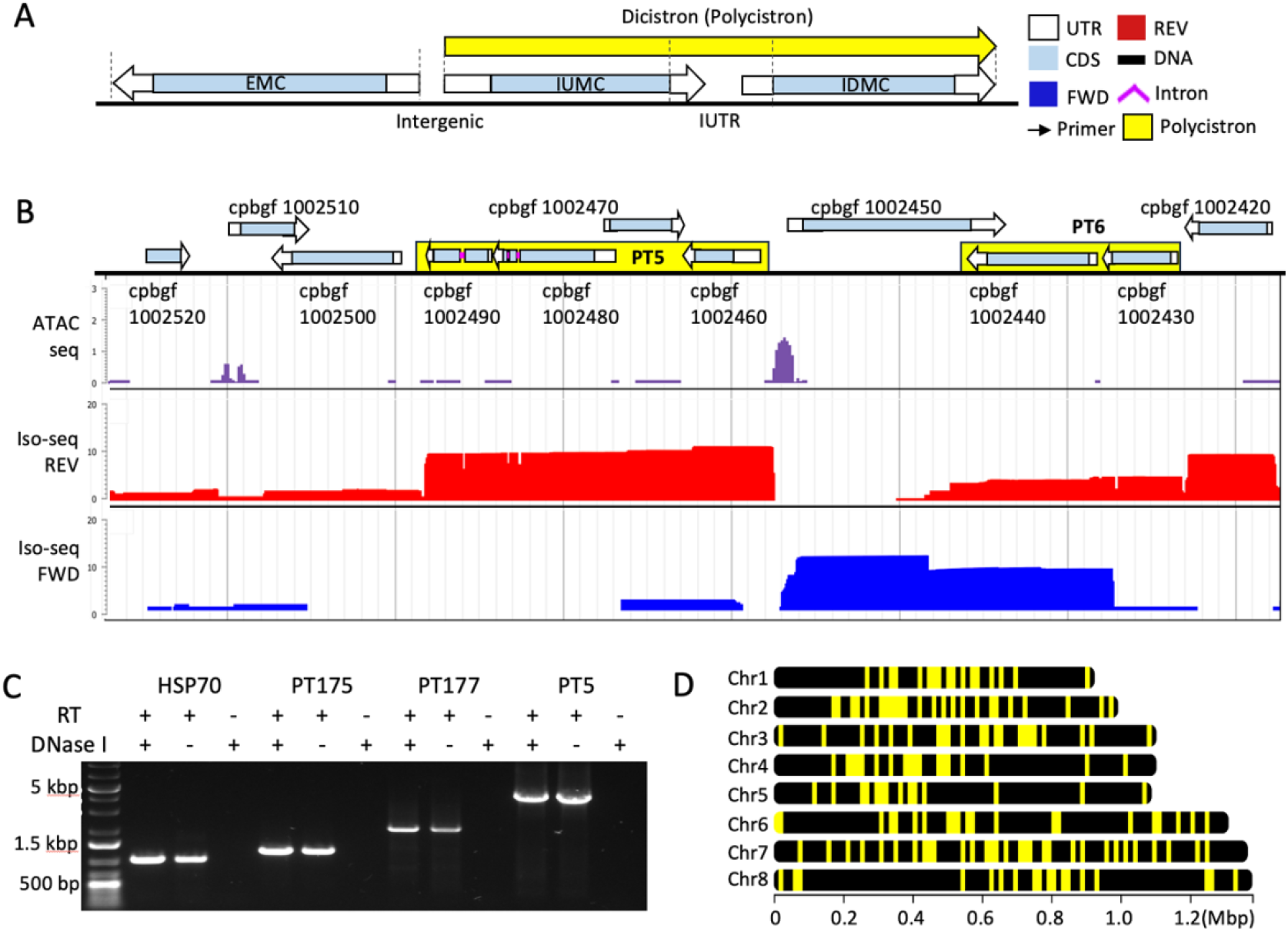
Discovery and validation of polycistronic transcripts in *C. parvum*. (A) Schematic representation of a polycistronic transcript (PT) showing an internal upstream monocistron (IUMC) and an internal downstream monocistron (IDMC) within a single PT transcript, alongside an external monocistron (EMC). (B) Transcriptional and epigenetic landscape for two polycistrons, PT5 (tricistron) and PT6 (discistron) on chromosome 1. Genes in PTs are highlighted in yellow boxes. ATAC-seq and Iso-seq tracks show reads mapped. ATAC-seq peak signal indicates accessible chromatin. Bigwig signals in all three tracks are transcript per million (TPM) normalized and log_2_ transformed. (C) RT-PCR validation of three select polycistrons, including two dicistrons (PT 175 and PT 177) and one tricistron (PT 5), with heat-shock protein 70 (*hsp70*) serving as a control. (D) Genomic localization of the 201 identified *C. parvum* sporozoite polycistrons.

## Results

### *Cryptosporidium parvum* expresses protein-encoding genes contained in polycistronic transcripts

The *C. parvum* IOWA-BGF haploid genome contains at least 9,259,183 bp spread across eight chromosomes and codes for 4,836 genes, of which 3,923 are protein-encoding(11). Full-length long-read RNA-seq revealed the presence of polycistronic transcripts within this compact, eukaryotic genome—a phenomenon typically observed in prokaryotes and certain eukaryotes. Unlike the canonical eukaryotic model, which supports a single ORF per messenger RNA (mRNA), our findings indicate that *Cryptosporidium* harbors multiple ORFs within single mRNA transcripts (Fig. 1). We illustrate the conceptual distinction between polycistronic transcripts (PTs) and typical monocistronic transcripts external to these polycistrons as external monocistrons (EMCs). We highlight the two unique genes internal to a dicistronic polycistron as the internal upstream monocistron (IUMC) and the internal downstream monocistron (IDMC), as shown in Fig. 1A. The regions between the PT’s internal genes are designated as the internal untranslated region (IUTR). PTs were detected from two distinct *Cryptosporidium* isolates with PacBio Iso-seq(30) and Oxford Nanopore (ONT) Direct RNA-Seq (DRS)(31) both of which are single-molecule long-read RNA-seq technologies (Figs. 1B, 2B). These results are not reverse transcription-based artifacts since the ONT DRS approach sequences native RNA molecules.

Mapping of single-molecule transcripts to the genome sequence also revealed that PTs have defined transcript boundaries (Figs. 1B, 2B, 3A). To further validate the PTs, RT-PCR was performed on three select polycistrons—two dicistrons (PT 175 and PT 177) and one tricistron (PT 5). Primers targeting the *hsp70* gene are used as a positive control(32). RT-PCR confirms that all three PTs are successfully amplified along with an *hsp70* control, even when DNase I is included (Fig. 1C). The 201 PTs identified in sporozoites cover all eight chromosomes (Fig. 1D). A Chi-squared analysis of PT genomic coordinates to assess random (null hypothesis) versus clustered distribution was inconclusive. The Chi-squared value of 9.6 is indicative of clustering but, the p-value of 0.2 is not statistically significant. PT chromosomal localization analysis supports a stochastic distribution rather than a clustered distribution, suggesting a non-selective genomic distribution of polycistrons. Overall, we conclude that polycistronic transcripts with defined boundaries exist in *C. parvum* sporozoites and these genes are not localized to a particular region of the genome. IDs, coordinates, types and internal monocistron gene IDs for all 201 polycistrons can be found in Supporting table 1.

### Polycistrons are validated using three different *C. parvum* biological sources

We were interested in examining the presence of PTs in additional *C. parvum* isolates. To further characterize and validate the existence of PTs, we examined long-read transcripts of an additional *C. parvum* IOWA isolate (Arizona) and a public post-infection (PI) 15 h timepoint from a different strain, INRAE(33). Taken together, the *C. parvum* PT landscape comprises 201 PTs, predominantly dicistrons (181), with the remainder being tricistrons (19) and a singular quadcistron. Interestingly there are two genes, cpbgf_8003620 and cpbgf_8003630 that occur as a Dicistron in the INRAE DRS data. Since we do not observe them as a PT with either BGF or Arizona isolate long-read RNA-seq data, and cannot use PCR to validate them, they are not included here. The 201 IOWA polycistrons contain 423 protein-encoding genes, representing a notable fraction (10.7%) of the organism’s protein-encoding gene repertoire (Fig. 2A). We discovered that PTs are also present in 15 h PI *in-vitro C. parvum* INRAE parasites (Fig. 2B). The results reveal the persistence of polycistronic transcripts into intracellular merozoite stage parasites, but there are some transcript differences related to developmental stage (Fig. 2B). Meta-profiling of PTs and EMCs across their transcript bodies using the three long-read RNA-seq datasets highlighted a consistent expression level and profile for PTs across their annotated transcript bodies (Fig. 2C). The Iso-seq reads for both PTs and EMCs show a slightly more prominent 5-prime expression bias versus the DRS reads, which have a slight 3-prime expression bias which may reflect the underlying differences between Iso-seq and DRS sequencing. The EMCs exhibit more variable expression levels across their transcript bodies between isolates, strains and developmental stage (Fig.2D) than the more similar PT expression levels (Fig. 2C). Additionally, the metaplot shows PTs have relatively well defined transcript boundaries, indicated by rapid expression level drops upstream and downstream of the TSS and TES for PTs similar to the transcript boundries observed in EMCs. These results show that PTs have defined boundaries and are not unique to *C. parvum* IOWA BGF isolate extracellular sporozoites, but rather, a more general phenomenon that extends beyond parasite strain and developmental stage.

**Fig 2.**
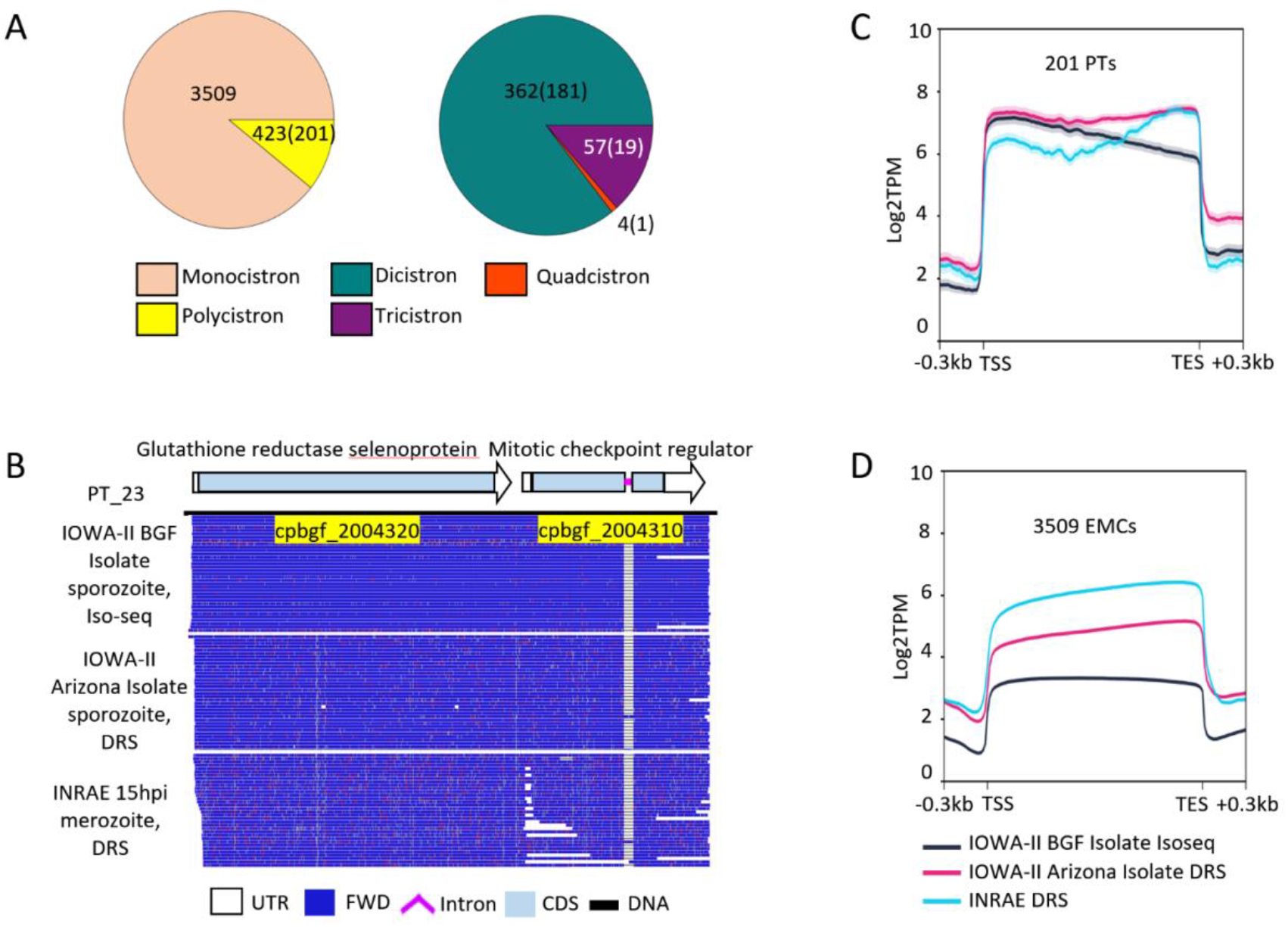
Abundance of polycistron types and expression profiles. (A) Abundance of transcript types. Numbers inside parentheses are PT counts, numbers outside are gene counts. (B) Long-read support of a dicistron from three different *C. parvum* sources. IMC IDs for PT genes are highlighted in yellow. Panels (C-D) Meta-profiling of expression levels for 201 PTs and 3509 EMCs. Expression signal is the mean log_2_(TPM). Colored lines represent the different isolates and strains as in the key. The line shadow indicates the standard error. Transcription start site (TSS) and transcription end site (TES) are shown along with upstream / downstream 0.3 kbp of the annotated regions.

To address whether PTs encode functionally related genes, we performed Gene Ontology (GO) enrichment analysis using the CryptoDB(34, 35) web service to compare PTs to EMCs for both molecular functions (MF) and Biological Process (BP). Results show only a subtle difference in GO term distribution was observed between PTs and EMCs, implying PTs do not encode a group of functionally distinct genes (Supporting table 3). PT IMC gene product descriptions were extracted from CryptoDB v68. IUMC and IDMCs within the same PT do not appear to share similar product descriptions, which suggests genes within PT share likely not of similar gene ontology terms (Supporting table 3).

We screened the *C. parvum* genome and sRNA transcriptome for trans-splicing leader sequences and did not find any significant hits. Homology-based searches for trans-spicing machinery also did not reveal any significant hits. These results suggest that the polycistrons found in *C. parvum* are likely not trans-spliced polycistrons.

### PT reads co-exist with IMC reads

Given that ORFs in PTs were annotated as independent genes and based on the observation of apparent internal monocistron transcripts, we set out to analyze the occurrence of internal monocistronic transcripts (upstream and downstream) present in annotated PT regions. To address this question, we performed an in-depth analysis of polycistronic transcription at the level of individual reads. Classified reads reveal the simultaneous presence of polycistronic and IMC reads. A genome browser view of this duality, featuring an internal upstream monocistron alongside their PT reads, is shown in Fig. 3A. We classified long reads based on their genomic alignment using minimap2(36) and the ORFs they encompass; those spanning the full length of annotated polycistrons were deemed PT reads, while reads encompassing a single complete ORF within polycistronic loci were categorized as either IUMC or IDMC. Since neither our Iso-seq nor DRS utilized 5′ guanine cap enrichment, reads that span incomplete ORFs are discarded as degraded transcripts. This approach minimizes potential false positives. A quantitative assessment of transcript abundance was conducted for all 201 identified PTs (Figs. 3B-D). We observe that most PTs have IMCs, with more abundant IUMCs than IDMCs. This pattern was preserved when evaluating the IOWA Arizona isolate reads obtained with DRS. However, this pattern is less prominent for the INRAE strain 15 h PI merozoites, along with a decrease in the overall abundance of PT reads. We performed an upset plot analysis to quantify these results (Fig. 3E). The results reveal that a large proportion (132 out of 201) of PTs contain both IUMCs and IDMCs alongside PT reads. When we compare IUMC versus IDMC presence, we observed that IUMCs and PTs are more prevalent compared to IDMCs. Interestingly, the higher the overall RNA expression, the more we see IMCs. These results suggest that PT transcription may be linked to local genomic transcriptional activity. In conclusion, the results show IMCs co-exist with PTs, which further adds to the complexity of this polycistron puzzle.

**Fig 3.**
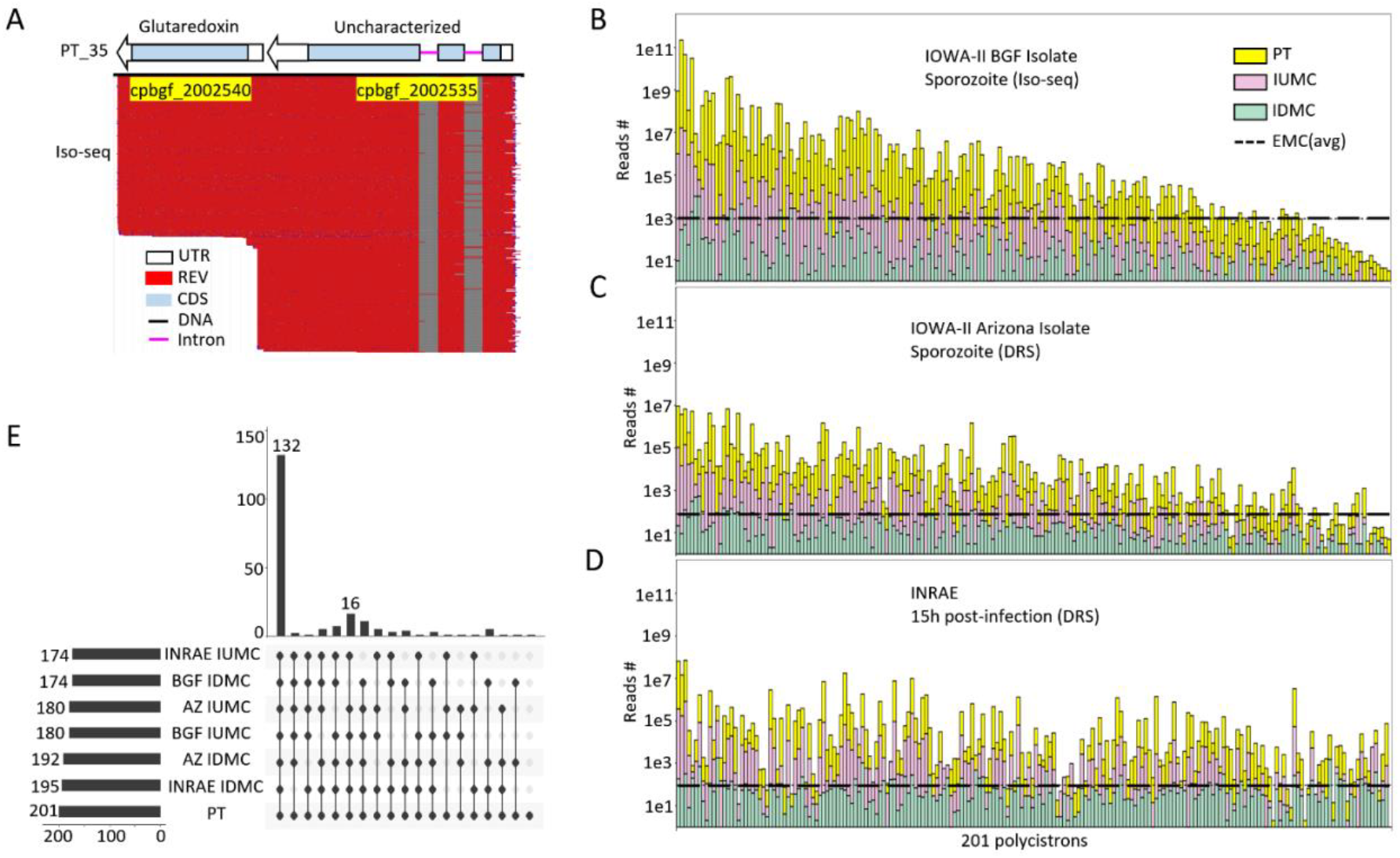
Internal monocistron transcripts co-exist with polycistron transcripts. (A) Genome browser view of an IUMC co-existing with PTs for PT35. (B-D) Read counts divided among PT, IUMC and IDMC for all three *C. parvum* long-read transcript sources, arranged from highest expression to lowest based on BGF Iso-seq on the x-axis. Log_10_ scaled raw read counts are used for the y-axis. The dashed line indicates the mean EMC read count per gene, from B to D at values of 864, 78, and 93 respectively. (E) Co-existence Upset plot of PTs, IUMCs and IDMCs for all three *C. parvum* RNA sources.

### ORFs inside PTs contain Kozak consensus sequences

We wanted to know if the ORFs encoded in PTs contained the requisite features to support translation into proteins. We looked for peptide evidence that could be mapped within polycistronic regions. *Cryptosporidium* has very limited proteomic data. We aligned peptide sequences to the genome via tblastn(37). In dicistrons, 10 of 181 have peptide support for both IUMCs and IDMCs, 32 of 181 only have IUMC peptide support, and 48 of 181 only have IDMC peptide support. In tricistrons and quadcistrons, 13 of the 20 have at least one IMC with peptide support. Taken together, 93 of the 201 PTs have peptide evidence. Fig. 4A, illustrates one dicistron example where both the IUMC and IDMC, which have different reading frames, have peptide support for both IMCs. We also found that PTs exhibit a greater proportion of peptide support compared to EMCs (Fig.4B). However, it is important to note that we do not know whether the peptide is translated using the PT as the template or the IMC(s). Analysis of the IUTRs within PTs and the intergenic regions between EMCs did not reveal any peptide mapping, thus bolstering the specificity of the findings (Fig. 4B).

**Fig 4.**
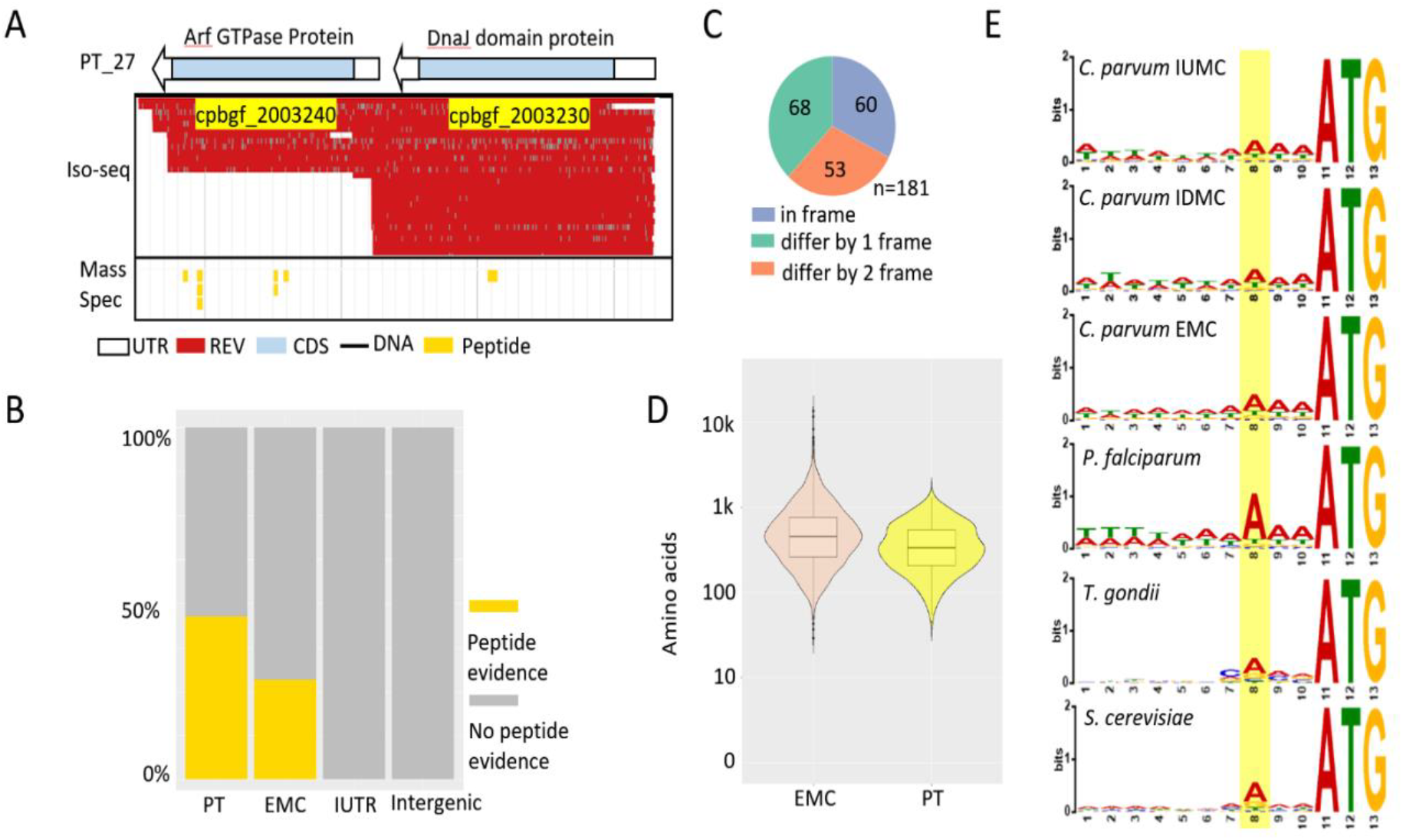
ORFs inside PTs contain evidence for translation of proteins. (A) Genome browser example of a dicistron PT27 with two ORFs not in frame. The top panel shows Iso-seq read support. The bottom panel shows mass-spec peptide support. (B) Peptide evidence comparison between PTs and EMCs. IUTRs and intergenic regions are used as a control. (C) Reading frame relationships for 181 dicistrons. (D) ORFs size comparison between EMCs and PTs. Units are of amino acids. (E) Kozak-like eukaryotic translation initiation sequence motif analysis between IUMCs, IDMCs and EMCs. *Plasmodium, Toxoplasma* and *Saccharomyces*’s annotated ORFs are used as an external control.

To address the potential issue of confounding misannotated genes with polycistrons, we analyzed the protein reading frames within the polycistrons. Focusing on dicistrons, the analysis revealed that only a third contain ORFs in the same reading frame (Fig. 4C). For the 19 tricistrons and the single quadcistron, four are in frame, while 16 have at least one ORF that is not in frame. The results suggest PTs are not mis-annotated monocistronic genes.

A comparison of the ORF size distribution between PTs and EMCs indicated no significant differences (Fig. 4D).

**Fig S1.**
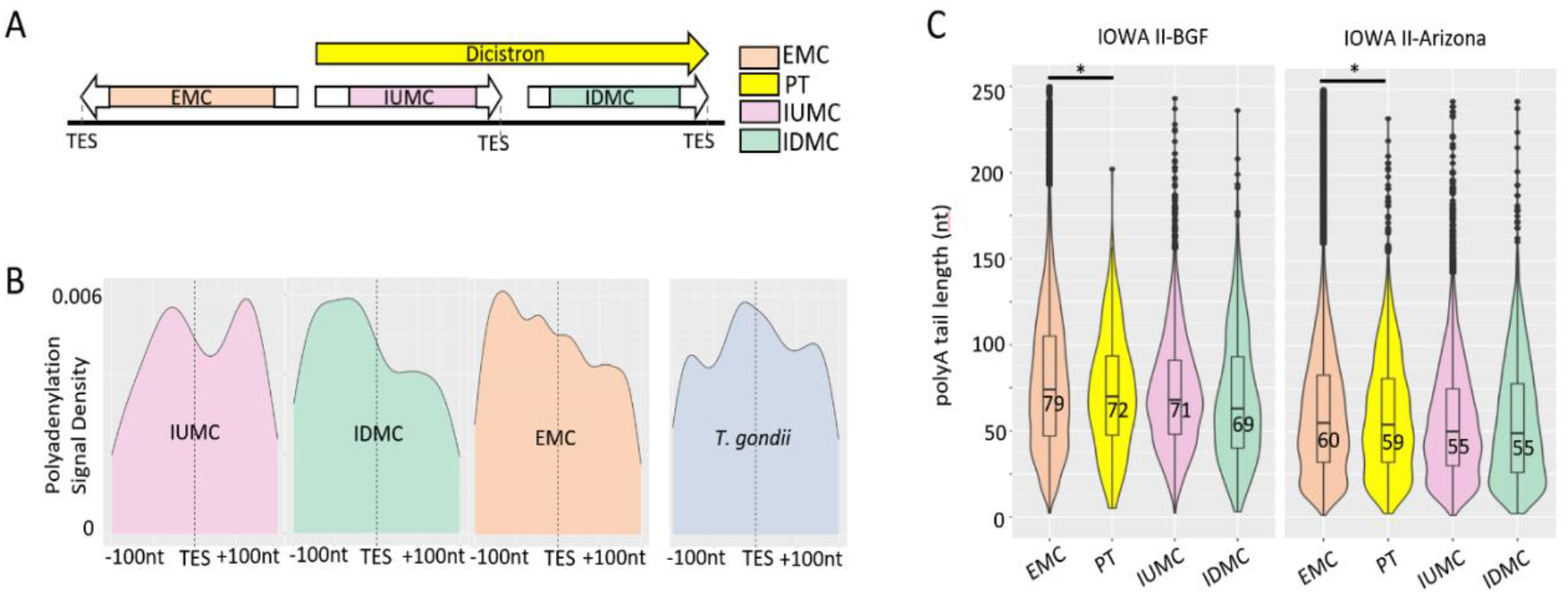
Polyadenylation signal motif and poly-A tail analysis. (A) Schematic of TESs among different transcript types. (B) Frequency density distribution of *Cryptosporidium* polyadenylation signals. The vertical dotted line marks the annotated TES. The area under the curve sums up to 100%. (C) DRS poly-A tail length distributions among different transcript types. Boxes represents 1^st^ to 3^rd^ quartiles. Line and number show average length. *p-value < 0.05 based on a two-tailed t test.

This observation was further supported by codon usage and amino acid usage comparison for ORFs within PTs and EMCs, which showed no remarkable variances (data not shown). Finally, the presence of Kozak-like sequences(38-40), crucial for translation initiation in eukaryotes, was confirmed in IUMCs, IDMCs, and EMCs. This motif was also compared to other apicomplexans(41) and yeast(42), establishing a similar propensity for translation initiation in PT cistrons (Fig. 4E). Both IMC and EMC’s codon usage as well as amino acid usage are calculated. In comparisons between IMCs and EMCs, we did not observe significant codon usage or amino acid composition differences (p-value =0.99, two-tailed pair-wise T test, Supporting Table 4). Thus, we conclude that ORFs encoded in PTs contain information sufficient for translation into proteins, while there does not appear to be major differences between PT and EMC regarding their codon or amino acid usage.

### Genes in PTs have proper polyadenylation signals, and PT reads have poly-A tails

To address the possibility that polycistrons are potentially generated from transcriptional read-through events, we interrogated PT, IMC, and EMC polyadenylation signals and poly-A tails. Polyadenylation signal sequences often signal transcription termination by RNA polymerase II in eukaryotic organisms(43). In *Cryptosporidium*, this motif is represented as 5′-A(A/G)TAAA(44). To assess the possibility that polycistrons result from the absence of polyadenylation signals in internal upstream monocistrons (IUMCs), we analyzed the distribution frequency of this motif around the transcription end sites (TES) for IUMCs, IDMCs, and EMCs, using the apicomplexan parasite *T. gondii* as a comparator. Results show that IUMCs and IDMCs exhibit polyadenylation signal profiles akin to EMCs, albeit with varied distribution profiles, none of which directly reflected the pattern observed in *T. gondii* (Fig. S1A-B). IUMCs have peaks of Poly-A signals before and after the TES site unlike IDMCs and EMCs which only have peaks before the TES.

To further address whether PTs are properly adenylated, we sought to analyze the poly-A tail length distribution for PTs. ONT’s DRS technology allows accurate measurement of poly-A tail length at a single molecule level(45). Utilizing the software suite Nanopolish(46), we directly measured poly-A tail lengths in individual DRS reads from the IOWA Arizona isolate. In addition, we conducted a supplementary low-depth DRS run on the same RNA sample used for the IOWA strain BGF isolate Iso-seq dataset to validate the poly-A tail length distribution findings. Our analysis confirmed the presence of poly-A tails on PT reads for both parasite sources. PT reads from both *C. parvum* sources have a tight distribution of poly-A tail lengths based on a quantile analysis. For both *C. parvum* sources, EMCs have statistically significantly longer poly-A tails compared to PTs, IUMCs, and IDMCs (Fig.S1C). No significant poly-A tail length differences exist among PTs, IUMCs, and IDMCs. Interestingly, we also observed that the *C. parvum* Arizona isolate has shorter poly-A tails overall versus BGF isolates. The results show that PTs are likely generated through proper transcriptional termination and poly-A addition.

### Not all polycistrons have accessible upstream regulatory regions in sporozoites, and very few have accessible IUTRs

As our data have shown that the majority of polycistrons have internal monocistron expression, we were curious whether IMC expression is driven by their own regulatory regions. To investigate the transcription initiation sites of *C. parvum* polycistrons, we conducted genome-wide chromatin accessibility profiling using ATAC-seq(47) on excysted sporozoites and we used genomic DNA (gDNA) from the same batch of excysted sporozoites as a control.

The genome browser visualization (Fig. 5A), and the insert size estimation (Fig. 5B) confirm the successful execution of ATAC-seq, specifically highlighting the tricistron PT5 with accessible chromatin exclusively at the 5′ upstream region of the annotated polycistron transcript model. PT5 does not have accessibility within either of its two IUTRs. PT5 does have detectable IMC expression of its first gene, cpbgf_1002460 in sporozoites. Focusing on dicistrons specifically (Fig. 5C), a comparative meta-analysis of dicistron accessibility versus EMCs revealed that most dicistrons present accessible chromatin only at their 5′ ends, akin to EMCs. When centering ATAC-seq signals on the IUTRs themselves, we see only a few dicistrons have accessible IUTR (Fig. 5C). This pattern held true for tricistrons and quadcistrons, each displaying prominent ATAC-seq signals at their 5’ upstream regions but not in their IUTRs (Fig.S2B).

**Fig 5.**
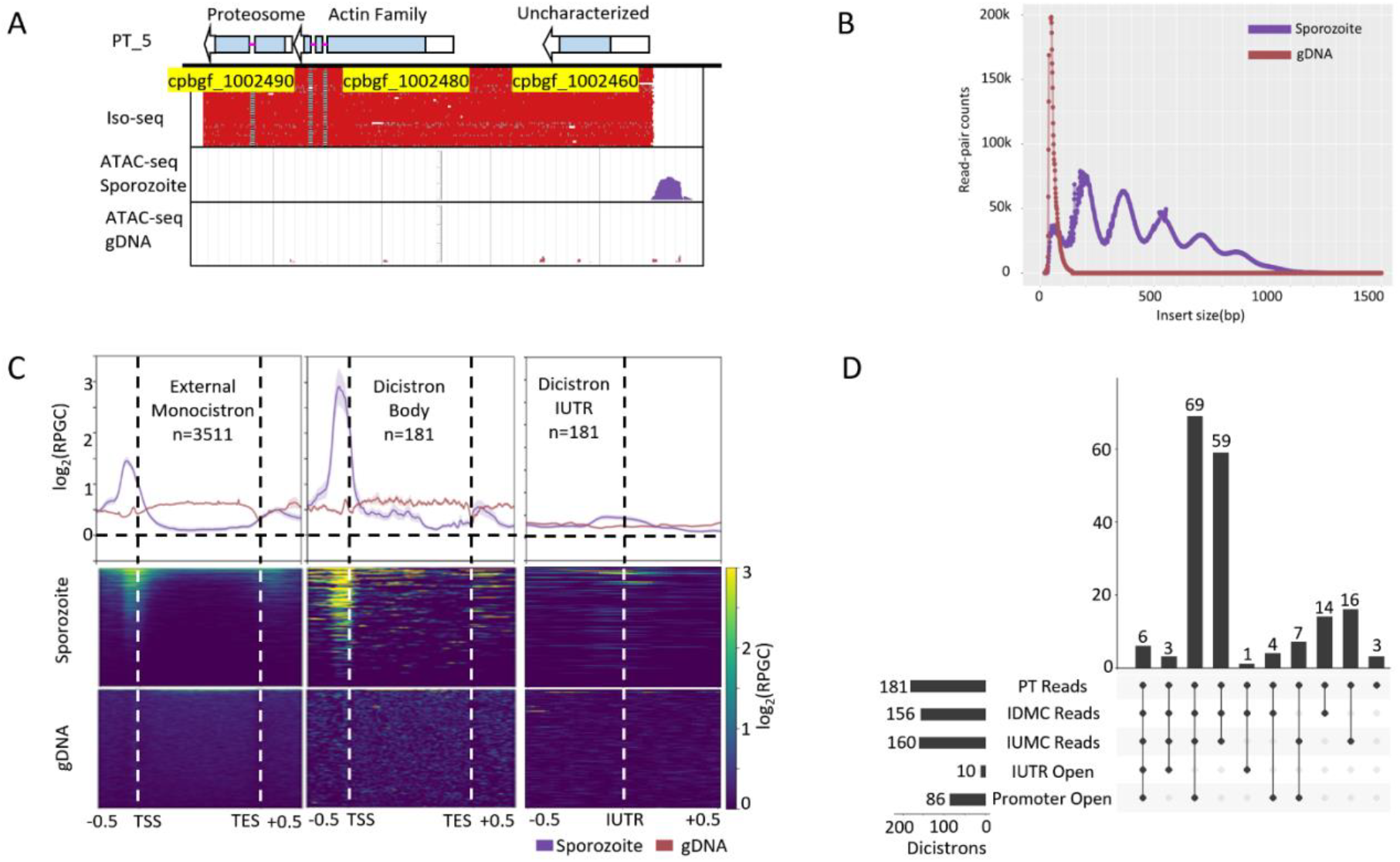
ATAC-seq analysis of polycistrons. (A) Genome browser of Iso-seq alignment and ATAC-seq signals for PT5. Sporozoite and gDNA ATAC-seq signals (shown in purple) are log_2_ transformed reads per genome coverage (RPGC). (B) Sporozoite and gDNA ATAC-seq insert size estimation distribution. (C) Meta-profile and heatmap analysis of EMCs (left), dicistrons (center) and dicistron IUTRs (right) chromatin accessibility landscape. Signals are log_2_(RPGC) transformed. (D) Integrative chromatin accessibility and long-read RNA-seq expression analysis in upset plot format.

**Fig S2.**
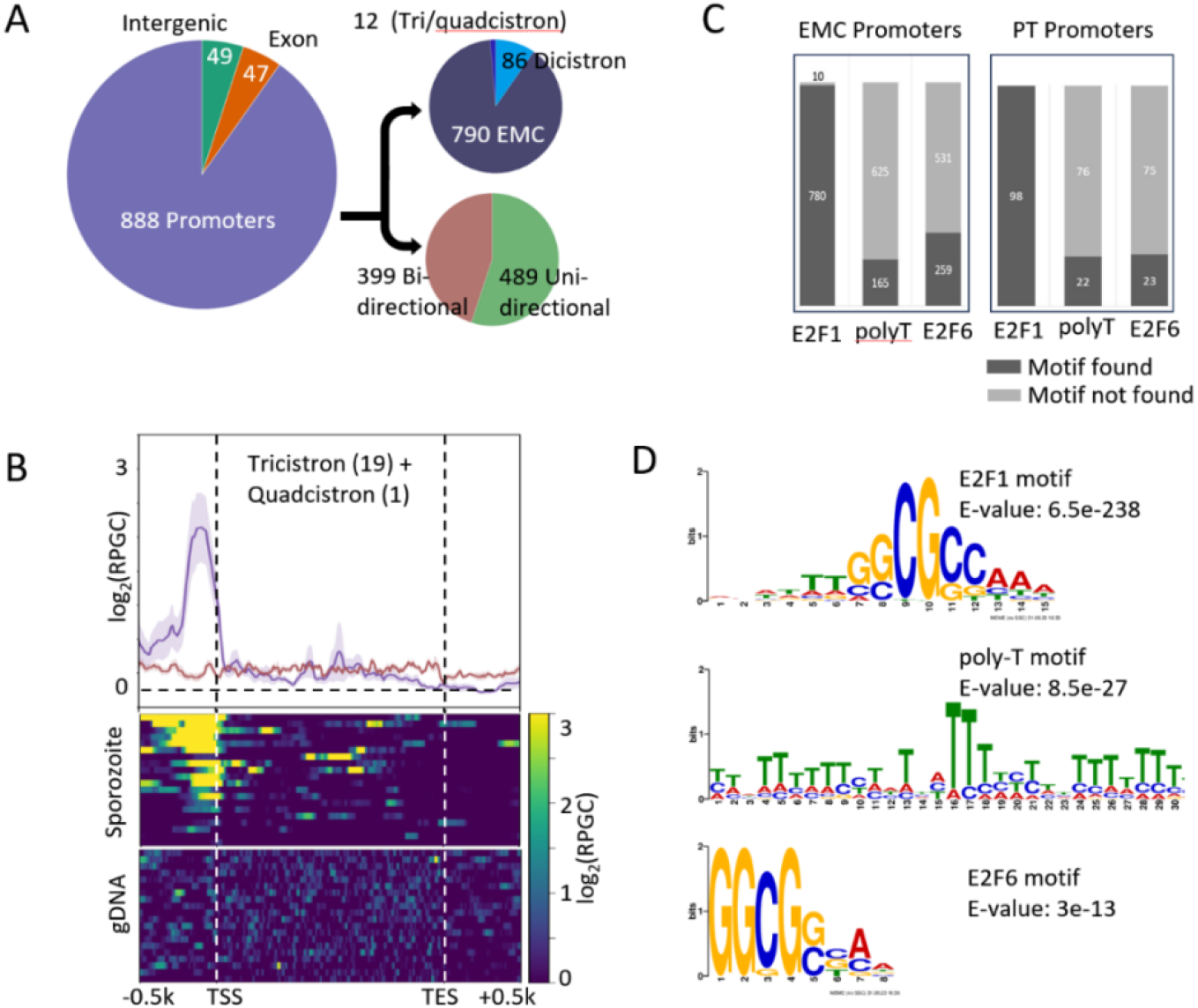
Upstream regulatory region annotation and motif analysis of PT ATAC-seq data. (A) Sporozoite ATAC-seq peak calling and annotation. (B) Tricistron and quadcistron chromatin accessibility meta profiling. (C) Annotated regulatory sequence motif composition comparison between PTs and EMCs. (D) Enriched regulatory motif logo and E-values using MEME suite.

The ATAC-seq profiling in sporozoites revealed that most sporozoite polycistron encoding regions do not appear to have genome accessibility around the PT 5′ end promoter region. To better understand this finding, we performed a quantitative integration analysis of Iso-seq and ATAC-seq data (Fig. 5D). The results are illuminating: 69 of 181 dicistrons possessed accessible PT promoter regions and PT, IUMC, and IDMC transcript reads but no accessible IUTRs to actively regulate expression of the IDMC. Another prevalent group included 59 dicistrons with PT, IUMC, and IDMC reads, but no accessible regulatory regions or IUTRs (Fig. 5D). These results demonstrate that IUTR accessibility does not correlate with the presence of internal downstream monocistronic transcripts and regulatory region accessibility does not correlate with PT transcript presence. Using the MACS3(48) HMMR model, we identified a total of 984 ATAC-seq peaks. The control data, which utilized genomic DNA, resulted in zero called ATAC-seq peaks. We annotated all 984 peaks with the HOMER(49) software suite. 888 peaks were categorized as upstream regulatory peaks, with most of them annotated as EMC regulatory regions, followed by dicistrons and a small number of tri and quad polycistrons (Fig.S2A). More than half of the regulatory peaks are annotated as bi-directional promoters (Fig.S2A). Sequence motif analysis using MEME suite(50) of ATAC-seq putative regulatory peak sequence regions revealed significant enrichment for E2F1 motifs, which were found to be abundantly present in both PT and EMC upstream regulatory peaks (Fig. S2C), indicating a shared regulatory feature. Motif enrichment analysis shows that E2F1 is the single most prominent motif identified, followed by a poly-T and the E2F6 motif (Fig. S2D). The poly-T motif has no known motif annotation using MEME suite’s Tomtom motif comparison tool(51). Interestingly, we do not observe any AP2 motifs in the detected ATAC-seq open chromatin regions. Results from this epigenetic profiling experiment have shown that IMCs are likely not driven by their own upstream regulatory regions in sporozoites. We have also scanned the IUTR sequences for any significant motif presence. Results show that the majority of PT IUTRs are enriched in E2F motifs (Fig. S3). Given the high frequency with which we discovered E2F motifs in ATAC-seq and IUTR regions, E2F transcription factors dominate gene regulation at the sporozoite stage and may play a role at other stages.

**Fig S3.**
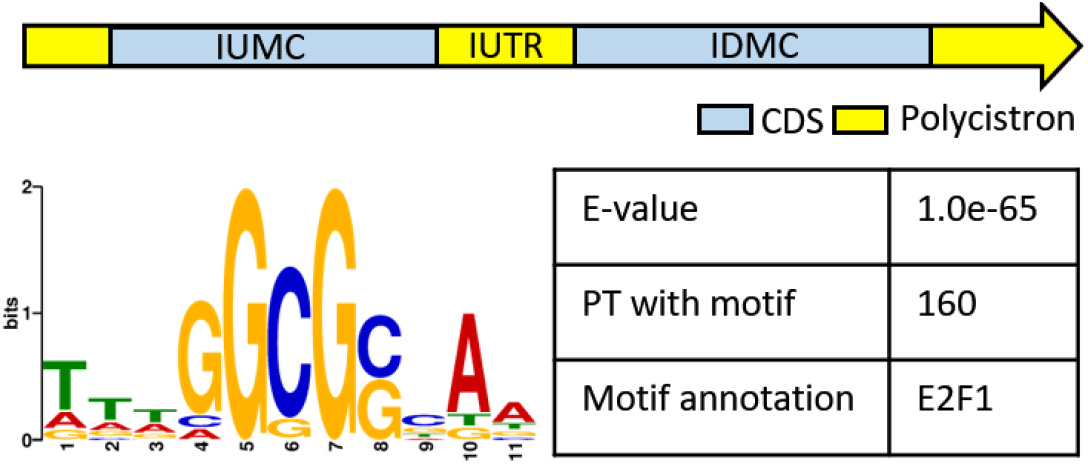
Sequence motif discovered at IUTR for all PTs. All 201 PTs’ IUTR regions are scanned with MEME motif discovery, allowing up to 10 motifs with default parameters. Only one motif shown here has E-value smaller than 0.05. 160 out of 201 PT’s IUTRs have this motif majority of PT IUTRs are enriched in E2F motifs (Fig. S3). Given the high frequency with which we discovered E2F motifs in ATAC-seq and IUTR regions, E2F transcription factors dominate gene regulation at the sporozoite stage and may play a role at other stages.

Given the high frequency of E2F motifs, we sought to utilize the recently published *C. parvum* single cell atlas to further characterize expression of the E2F and DP genes across the life cycle(52). There are two annotated E2F genes in the *C. parvum* genome: cpbgf_1001560 and cpbgf_6001430. In addition, there are two DP genes (cpbgf_7003650 and cpbgf_8001850) which encode co-regulators of E2F genes. Mapping individual E2F/DP gene expression onto the single cell atlas UMAP reveals that both E2F and DP are highly enriched in asexual stages (Figs. S4A-D). Further binning of E2F and DP gene expression across the 18 previously defined life stage clusters shows that the E2F and DP genes are highly expressed early in parasite development, clusters 1 and 2. They also have moderate expression late in the asexual stage, clusters 8 and 9 (Fig. S4E). We observe a small number of cells expressing both E2F and DP, as well as lower expression levels in developing male and female gamont clusters. In female gamonts we see low levels of E2F and DP expression, and an even lower level of expression in male specific clusters. These results show that E2F, and DP are mainly, but not exclusively, expressed during asexual parasite stages thus likely contributing to asexual gene regulation.

### Some *Cryptosporidium* PT expression profiles change with development and do not appear to be enriched in either male or female restricted genes

Most PTs have co-expressed IMCs, usually the IUMC. Using short-read RNA-seq data, we also observe IMC only expression appearing later in development. Without sufficient long-read RNA-seq evidence for merozoite and gametocyte stages we cannot determine proper IMC boundaries, assuming there are boundaries at all. In fact, we do not know if the IMCs represent *de novo* transcripts or if they are generated via processing of the PT. We utilized available short-read RNA-seq data to examine gene expression at polycistronic loci in merozoites. Looking specifically at PT198 (Fig. 6A), PT177 (Fig. 6B) and PT92 (Fig. 6C) at the 0 h (sporozoite) time point, we observe short-read support for the polycistron annotation, with RNA-seq reads continuously aligned across PT region. However, when we examine the 24 h time point replicates, the expression pattern changes dramatically for all three dicistrons. At the 24 h time point, PT198 has completely lost its IDMC expression and IUMC expression is reduced compared to the 0 h time point. PT177 and PT92 each have lost PT expression, as evidenced by the loss of continuously mapped short-read RNA-seq data across the IUTR junction. PT177 has a large intron. Interestingly, there are some intron mapped reads at the 24 h time point while all IUTR mapped reads are lost. Sporozoite chromatin accessibility revealed by ATAC-seq for these PTs shows open chromatin at all upstream PT promoter regions and PT177 showed IUTR accessibility similar to its IUMC promoter region, likely explaining the co-existence of PT177 and its IDMC transcripts. Although we cannot use bulk short-read RNA-seq to quantitatively determine polycistron differential expression patterns across developmental stages, our qualitative analysis does reveal changing polycistron and IMC transcriptional profiles during development.

**Fig S4.**
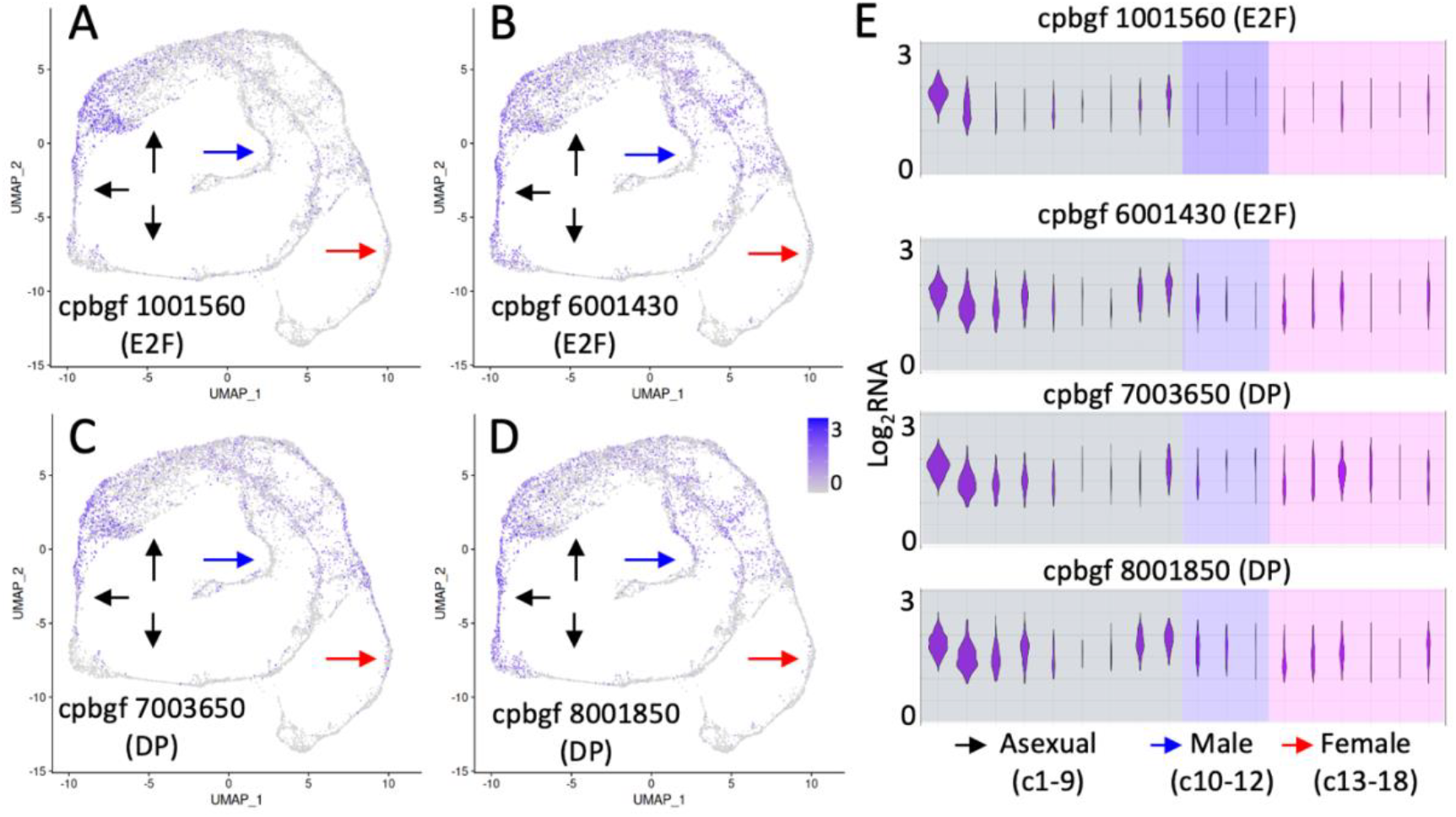
Single cell expression analysis of E2F and DP genes. (A-D) UMAP expression mapping to single cell atlas for E2F and DP genes. Purple highlights cells with high expression, while grey indicates cells with no E2F expression. Expression levels of E2F and DP are scaled from grey to purple using log_2_ transformed RNA expression. Arrow indicates asexual, male and female stages. (E) E2F and DP expression level aggregated by individual cell clusters arranged in asexual, male and female specific gene clusters. Violin plot area positively correlate with number of cells with observed gene expression. C is short for cluster.

**Fig 6.**
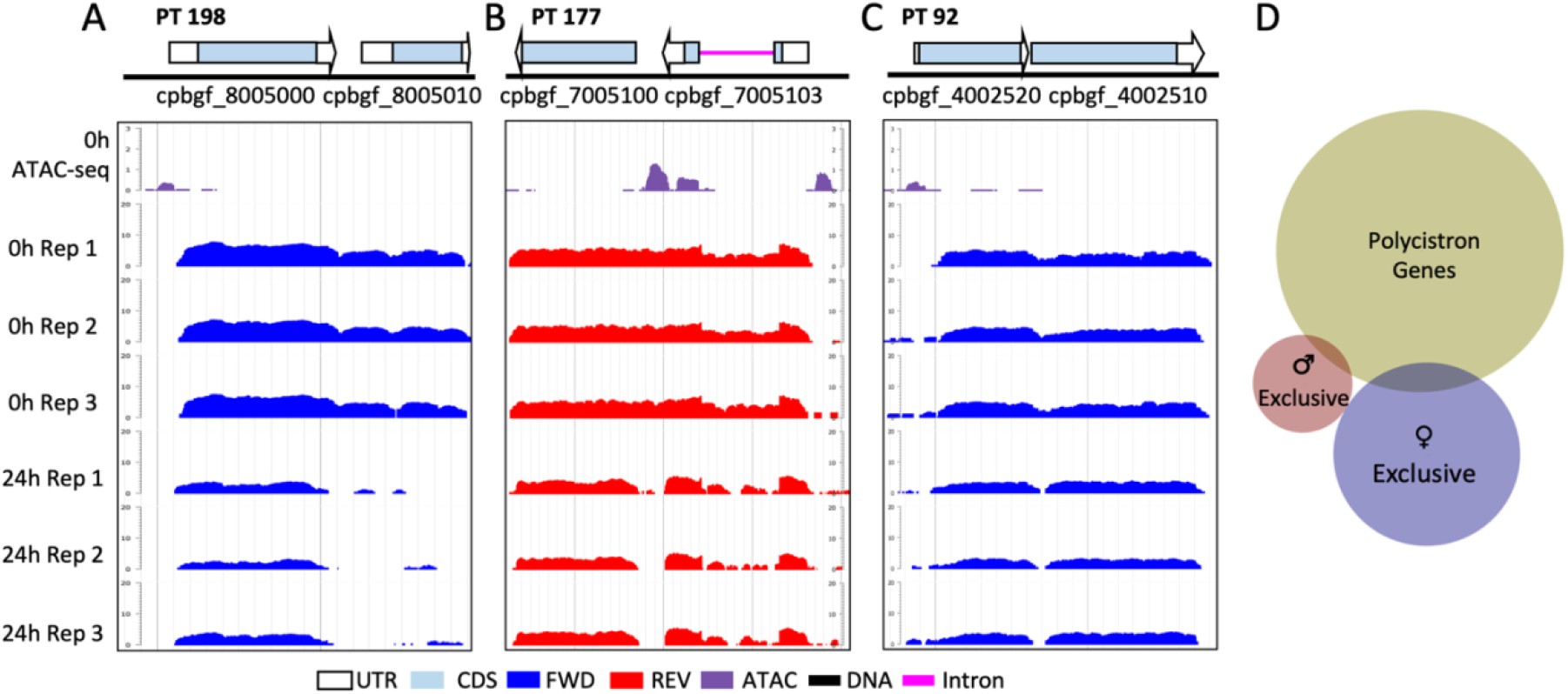
Developmental differences in dicistronic RNA presence. (A) PT198 (B) PT 177 and (C) PT 92 with their RNA expression in triplicate per time point, and sporozoite chromatin landscape. Short-read RNA-seq and ATAC-seq coverage is shown in bigwig format. Per base-pair coverages are normalized with TPM and log_2_ transformed so bigwig scores can be compared directly. cpbgf_8005000: utp12 domain-containing protein. cpbgf_8005010: uncharacterized protein. cpbgf_7005100: thioredoxin. cpbgf_7005103: uncharacterized protein. cpbgf_4002520: uncharacterized protein. cpbgf_4002510: uncharacterized protein. (D) Venn Diagram of PT, male exclusive and female exclusive genes. PT: 423 genes, male () exclusive: 51 genes, female () exclusive: 187 genes.

**Fig S5.**
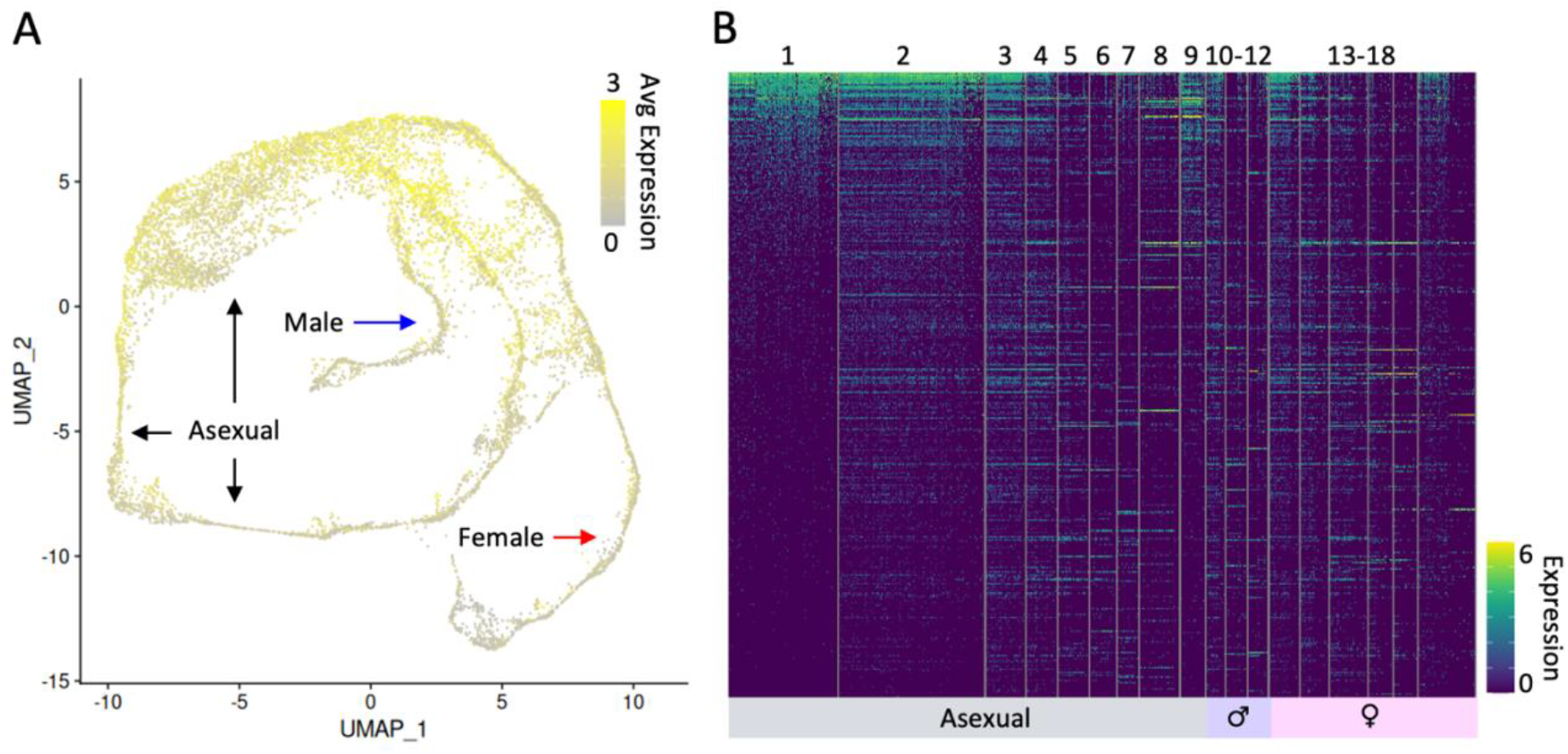
Expression analysis of 423 PT encoded genes in the *C. parvum* single-cell atlas. (A) UMAP of aggregated average PT gene expression mapped to *C. parvum* cells across developmental stages. Color scale represents mean PT gene expression for individual cells. Arrow indicates asexual, male and female stages. (B) PT gene expression heatmap across all 18 previously annotated cell clusters from three developmental stages. Each row represents one PT encoded gene, each column represents one cell. Cluster width indicates number of cells with detected RNA expression for a given PT gene.

Analysis of *C. parvum* single-cell RNA-seq has revealed genes that are specific to either male or female development (52). We used the *C. parvum* scRNA atlas to analyze PT gene expression patterns. Aggregating the mean expression of all 423 PT genes together reveals that PT genes are expressed across the entire *C. parvum* life cycle. Despite this finding, the UMAP does show that PT genes are highly expressed during asexual stages (Fig. S5A). We also mapped expression of all PT genes to the defined 18 gene clusters from the *C. parvum* scRNA atlas. Heatmap results show cells expressing PT genes are mostly found during asexual stages indicated by the large column width of asexual stage clusters, especially clusters 1 and 2 (Fig. S5B). Comparing polycistron genes to genes expressed exclusively in developing male and female gamonts, we found that 404 of 423 PT genes are not exclusive to (Fig.6D). We found 4 male exclusive genes that are also present in PTs, cpbgf_2002860, cpbgf_3003550, cpbgf_5001830 and cpbgf_8002980. Among them, cpbgf_5001830 is functionally predicted to be a zinc-finger containing protein which suggests it may be a transcription factor. We also found 15 female exclusive genes that are present in PTs, cpbgf_2001220, cpbgf_2001590, cpbgf_3001860, cpbgf_3002870, cpbgf_4003090, cpbgf_5001210, cpbgf_5001370, cpbgf_7002990, cpbgf_7003120, cpbgf_7003760, cpbgf_7004890, cpbgf_8002360, cpbgf_8002370, cpbgf_8002670 and cpbgf_8003133. 5 out of these 15 genes are annotated as uncharacterized proteins, and the rest are functionally diverse and do not appear to enrich to any specific set of GO terms. This result suggests that majority of genes in PTs are likely not linked to *C. parvum* ‘s sexual development.

## Discussion

The robustness of our methodology and the consistent detection of polycistrons across independent *Cryptosporidium* isolates and sequencing technologies negate the likelihood of PTs being mere transcriptional anomalies. The precise transcript boundaries and the RT-PCR validations reinforce the authenticity of the findings. Eukaryotic polycistronic transcripts typically adhere to one of two models: *trans*-splicing, as in trypanosomes, where a leader RNA sequence is appended to facilitate mRNA maturation and subsequent translation, resulting in monocistronic mRNAs, or non-*trans*-splicing, observed in certain algae, where a subset of genes is polycistronic with inherent *cis* splicing. Protein sequence homology-based analysis could not identify any *trans*-splicing machinery genes in *C. parvum*. In addition, we scanned the *C. parvum* genome for potential *trans*-splicing leader sequences using trans-splicing detection tool, SLIDR-SLOPPR(53). The tool did not detect any *trans*-splicing leader sequences present in publicly available RNA-Seq datasets, further suggesting that the polycistrons in *C. parvum* are not *trans*-spliced. Furthermore, our lab has previously conducted small RNA-seq on *C. parvum*(16), and none of the novel annotated small RNA genes were characteristic of spliced-leader sequences, which range in length from 15 to 50 nt, are present in high abundance, and map to 5’ end of transcripts(54). Given the absence of *trans*-splicing machinery genes and *trans*-splicing leader sequences, coupled with a comprehensive *cis*-splicing system(55, 56), our data align with the latter model. Thus, *Cryptosporidium* polycistrons seem akin to those detected in red and green algae, functioning without *trans*-splicing. The predominance of dicistrons within *Cryptosporidium*’s polycistronic profile aligns with green and red algae observations, diverging from the *trans*-splicing polycistron paradigm predominant in organisms like trypanosomes.

The stable expression patterns and defined transcriptional boundaries of polycistrons contrast with the more variable expression of EMCs, suggesting a distinct regulatory framework for PTs. The overall similarity in GO term composition between polycistrons and EMCs may reflect a shared biological imperative, with polycistrons potentially mirroring the functional diversity of the broader gene pool as shown in the GO term comparisons between PTs and EMCs (Supporting table 3). We compared genes present in PTs to the list of 190 genes that are determined to be under positive selection(10). Results show none of the PT genes are present in the list of genes under positive selection. This suggests that PT gene functions are likely not directly related to interactions with the host.

The simultaneous presence of PTs and IMCs within *C. parvum* suggests a complex gene regulatory mechanism that likely involves both transcript initiation and transcript termination. For an organism that relies on salvaging host nucleotides for DNA and RNA synthesis, the co-production of PTs and IMCs might seem energetically extravagant. However, our observations imply that the advantages of this transcriptional arrangement may supersede the apparent resource expenditure. The presence of IUMC and IDMC reads, while variable, is typically subordinate (they are less abundant) to PTs (Fig. 3B-C) in sporozoites. In contrast, the INRAE strain, which reflects the parasite’s post-infection merozoite transcriptomic profile, displays a more balanced representation of PTs and IMCs. Notably, the relative abundance of IUMs approaches that of PTs, suggesting a shift in gene expression regulation during the intracellular life cycle stage (Fig.3D). Given the lack of single molecule long-read RNA-seq data for later asexual stages as well as sexual stages, we cannot determine whether PTs are present across the entire life cycle of the parasite. However, leveraging publicly available single-cell RNA-seq data we gained some insight into expression of the genes contained in PTs across the full life cycle. These data show that genes encoded in PTs are indeed expressed across all life cycle stages but with a bias towards asexual stages. It is plausible that PTs are made along with IMCs post fertilization during new oocyst maturation. This could explain the large number of PT transcripts that were not associated with open chromatin in sporozoites. The abundant number of coding transcripts (IMC + PT) may allow sporozoites to quickly start their development upon infecting host cells. This is when E2F and DP are abundantly expressed. But, we do not know if all ORFs contained within the PTs are translated.

The detection of peptide evidence from available mass spectrometry data confirms the translational activity of ORFs within PTs, particularly when contrasted with the lack of such evidence within IUTRs for PTs and intergenic regions outside of PT locus. Having ribosomes continue translation after hitting stop codons is difficult to imagine without confirmed IRES sites. The similarity in ORF sizes and codon usage between PTs and EMCs in addition to their annotated functions (when known), suggests that the ORFs coded by PTs likely produce proteins. Previous research has shown that when the distance between two ORFs is relatively close, ribosomes may be able to reinitiate translation through leakage scanning of the transcripts until a new translation initiation Kozak-like sequence motif is found(38). Research in green algae polycistrons has already demonstrated that such a mechanism can translate two proteins from a single transcript *in vitro*. Research in other apicomplexan parasites, such as *Plasmodium falciparum* and *Toxoplasma gondii*’s upstream open reading frames (uORFs), has also demonstrated the existence of ribosome leaky scanning and reinitiation(41, 57-59). The identification of Kozak-like sequences in both IUMCs and IDMCs, comparable to EMCs, reinforces the potential for the PT ORFs to be translated. This, coupled with the absence of significant disparities in amino acid usage and tRNA codon usage (Supporting table 4), suggests that the translational machinery may recognize ORFs inside PTs similarly to ORFs inside EMCs, enabling diverse protein expression from a single polycistronic transcript. Translation of multiple peptides from a single transcript may offer higher translation efficiency(60), which can be beneficial for an intestinal parasite under host immune pressure to invade host epithelial cells successfully. These observations underscore the complex nature of *C. parvum* PTs and suggest that they are an integrated, functional component of the parasite’s gene expression machinery.

The presence of polyadenylation signals and poly-A tails for IUMCs suggests that transcription can be terminated at IUMCs(61). IUMCs, IDMCs and EMCs share a similar polyadenylation signal profile, indicating the presence of canonical polyadenylation signals across different transcript types. Poly-A tail length distributions among PTs, IUMCs and IDMCs are comparable. Previous research has shown an inverse correlation between poly-A tail length and translational efficiency(62). The notably shorter poly-A tails in PTs compared to EMCs could indicate a higher translation efficiency for PTs, as previously reported in *C. elegans*(62).

Epigenomic research is lacking for *Cryptosporidium*. Only recently has research been performed to validate the existence of functional histone modifiers in the parasite via enzyme activity assays and certain histone marks via immunomicroscopy(63). The ATAC-seq(47) analysis offers a revealing glimpse into the epigenetic regulatory landscape in *C. parvum* sporozoites. The nucleosome spacing periodicity of ATAC-seq signals underscores a robust chromatin accessibility pattern. We observed a substantial number of PTs, as well as EMCs, that lack accessible upstream regulatory regions despite the presence of ample RNA transcripts. This suggests that transcriptional activity may be limited in sporozoites, inside oocysts, waiting and poised for excystation and infection. Some PTs, as well as EMCs, were likely transcribed or deposited during sporozoite formation during oocyst development.

The predominance of E2F motifs within identified ATAC-seq peaks is particularly intriguing. Like other apicomplexan parasites(64), *Cryptosporidium* has a reduced repertoire of transcription factors, and relies heavily on a family of AP2 TFs(65). The absence of AP2 DNA binding motifs in sporozoite open chromatin regions is notable. Thus far, *Cryptosporidium* appears to be the only genus within the phylum Apicomplexa where E2F transcription factors have been retained(66). Drawing parallels from mammalian studies of E2F transcription factors, which function in cell cycle regulation(67-69), it is conceivable that *Cryptosporidium* may utilize E2F-driven transcription for host invasion and asexual reproduction. The *C. parvum* genome has two annotated E2F transcription factor genes and two DP genes, which are E2F binding partners. Interestingly, we have found that one of the E2Fs, cpbgf_7003650, resides within a dicistron, PT168. The E2F and DP genes themselves are apparently driven by upstream regulatory regions with E2F binding motifs. This suggests that potentially E2Fs may autoregulate(70). Sexual differentiation in *C. parvum* is hypothesized to be heavily regulated through AP2 gene families(8, 52, 65). Our analysis of genes encoded in PTs suggests their expression is not enriched in developing gamonts. Given our observation of the overwhelming presence of putative E2F binding sites in open chromatin in sporozoites, upstream of PT genes and between IMCs in PTs, suggests that PT genes and early development are likely controlled by E2F regulated transcription.

Our investigation of polycistrons relied heavily on data derived from the parasite’s sporozoite stage, especially the long-read RNA-seq datasets. This approach inherently introduces a bias, potentially overlooking the scope of polycistronic transcriptional activity across various developmental stages. Furthermore, the employed long-read RNA-seq does not discriminate based on the presence of the 5’ guanine caps in RNA molecules, which complicates our ability to discern the co-occurrence of PTs and IMCs. Without explicit TSS annotation, such as R2C2(71) or Tera-seq(72), we cannot conclude whether IMCs are dependent on processing of PT transcripts (Fig. S6A), or are independently transcribed using their own promoters (Fig. S6B). Although ATAC-seq reveals open chromatin regions, our analysis does not extend to identifying specific histone modifications flanking these accessible areas. Despite the presence of poly-A tails on PTs, IUMCs, and IDMCs, the absence of data on transcription termination and histone modifications, such as H3K36me3 enrichment, makes it difficult to determine whether the production of PTs is due to a failure in transcription termination. Such limitations render it difficult to study transcription factor binding and epigenetic regulations of polycistrons. Single-cell RNA-seq based atlas studies are powerful in differentiating *C. parvum* cell types across developmental stages. Analysis of *C. parvum* single-cell RNA atlas data supports our hypothesis that PTs are important during early asexual development. However, due to the sparsity of single-cell RNA-seq data(73), and the inherent limitations of short-reads used in this approach, we do not know whether the expression of genes located in PTs comes from IMCs or PTs in these later life cycle stages. Our study, though revealing, represents an initial investigation into the complex RNA biology in *C. parvum*, highlighting the need for further detailed studies.

**Fig S6.**
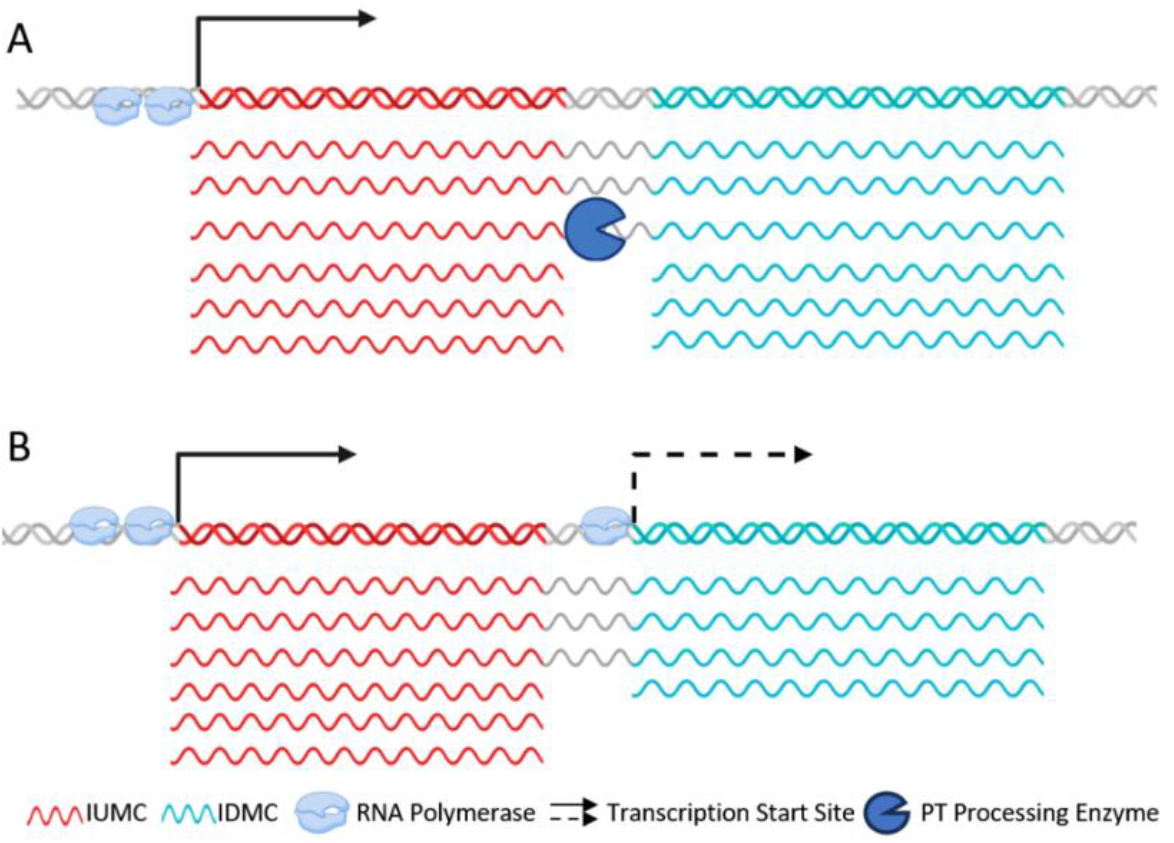
Proposed models of IMC origin. (A) Dicistron is transcribed as a whole with both IUMC and IDMC, thus the IUMC does not terminate properly. Separate IMC molecules are then generated from processing of polycistron RNA molecules. (B) Dicistrons and IMCs are independently transcribed. As in model A PTs are generated by failure of the IUMC to terminate properly. The Dashed line represents less accessible promoters. RNA polymerase numbers represent local transcription activity.

The revelation of polycistronic transcripts within the *C. parvum* genome significantly advances our understanding of the parasite’s gene regulation. This study confirms the presence of polycistronic transcription in a member of the Apicomplexa and delineates the regulatory complexity of these transcripts while highlighting the importance of E2F-based gene regulation. Our comprehensive analysis, leveraging the synergy of long-read RNA-seq, ATAC-seq, short-read RNA-seq, and single-cell RNA-seq has established that polycistrons in *C. parvum* are processed like monocistronic messages, complete with poly-A tails and developmental stage-specific expression. In summary, our findings challenge the long-standing paradigm that polycistronic mRNAs are rare in eukaryotes. Our data suggest that polycistronic transcription is a regulated, potentially functional, and evolutionarily conserved mechanism within *Cryptosporidium*. Additionally, we hypothesize that polycistrons may have arisen in *C. parvum* due to its compact 9.2 Mb genome size relative to most other apicomplexan parasites. The smaller genome limits the available intergenic regions for *cis*-regulatory elements. Consequently, with a diminished repository of *cis*-regulatory elements, two or more genes can be regulated and expressed together within a single transcript, forming a polycistron. These insights broaden our understanding of the molecular biology of *C. parvum*. This study paves the way for future research into the prevalence and significance of polycistronic transcripts in other *Cryptosporidium* species and potentially other apicomplexan parasites.

## Materials and Methods

### *C. parvum* sporozoite cell prep

*C. parvum* IOWA-II strain oocysts were obtained from Bunch Grass Farm (BGF) (Deary, ID) and the *Cryptosporidium* Production Laboratory at the University of Arizona (Tucson, AZ). 1.0×0^8^ oocysts were resuspended in cold Dulbecco’s phosphate-buffered saline (DPBS) and incubated in fresh household bleach (20% mixed in DPBS) on ice for 10 mins and then washed twice with DPBS. Excystation of sporozoites was induced by incubating oocysts in 0.8 % sodium taurodeoxycholate (Sigma) in DPBS at 15 °C for 30 mins, then 37 °C for 1 h as previously described(15). Excysted sporozoites were briefly centrifuged to collect them. The sodium taurodeoxycholate supernatant was removed, and sporozoites were resuspended in cold DPBS for downstream usage. Excystation was verified through imaging and counting with Cellometer X2 (Nexcelom).

### *C. parvum* IOWA-II strain BGF isolate RNA isolation and sequencing

Approximately 4.0×0^8^ excysted sporozoites from Bunch Grass Farm were incubated with 1 ml of RLT buffer with 10 ′l of β-mercaptoethanol (Sigma-Aldrich) added. RNA extraction was performed using the RNeasy Mini Kit following the “Purification of Total RNA from Animal Cells” protocol from the manufacturer (QIAGEN). Extracted RNA was treated with DNase I (Invitrogen) to remove DNA contamination. Cleaned RNA was resuspended in nuclease-free water and quantified using a TapeStation (Agilent). 1 ′g of total RNA was used for input into PacBio Iso-seq library construction. The prepped library was sequenced on a Sequel II system at the Georgia Genomics and Bioinformatics Core. Additionally, 50 ng of total RNA from the Iso-seq source was input into ONT DRS library prep and sequenced on a R9.4.1 flowcell for 24 h.

### *C. parvum* IOWA-II strain Arizona isolate RNA isolation and sequencing

Approximately 4.0×0^8^ excysted sporozoite cells from the *Cryptosporidium* Production Laboratory at the University of Arizona were incubated with 1 ml of TRIzol (Invitrogen) and incubated at room temperature with gentle shaking for 10 mins. Total RNA isolation was performed following the TRIzol protocol. Extracted RNA was treated with DNase I (Invitrogen) to remove DNA contaminations. Cleaned RNA was resuspended in nuclease-free water and quantified using a Nanodrop. 1 ′g of total RNA was used for input into an Oxford Nanopore Direct RNA-sequencing library kit, RNA002, following the manufacturer’s protocol. The library was loaded onto an ONT R9.4.1 FLO-MIN106D flowcell, and sequenced in a GridION for 72 h.

### *C. parvum* ATAC-seq tagmentation and sequencing

Approximately 4.0×0^7^ excysted sporozoites in DPBS were centrifuged at 16,000g at 4 °C for 3 mins. The pelleted sporozoites were gently resuspended in 22.5 ′l of 0.1% digitonin (in DPBS) solution. Cells were incubated on ice for 3 mins to permeabilize the cell membrane. 25 ′l of Tn5 reaction buffer and 2.5 ′l of TDE1 Tn5 adapter-bound tagmentation enzyme (Illumina) were added to the permeabilized cell lysate. The reaction was incubated at 37 °C for 45 mins, with gentle flicking every 10 minutes. As a control, genomic DNA was extracted from the same excysted sporozoites with phenol:chloroform:isoamyl alcohol (Invitrogen) following the manufacturer’s protocol. Genomic DNA was treated with RNase A to remove RNA contamination and cleaned up with the Blood and tissue kit (QIAGEN). 1 ng of genomic DNA was used for tagmentation as a control. The *C. parvum* IOWA-II strain BGF isolate tagmentation products and tagged gDNA control were purified using a Monarch PCR & DNA Cleanup kit (T1030S, NEB) and then amplified using NEBNext® High-Fidelity 2X PCR Master Mix (M0541S) for 11 cycles following the manufacture’s protocol. Amplified libraries were purified with AMPure XP beads, and library concentrations were determined using a Qubit and qPCR prior to sequencing. Libraries were sequenced with Illumina NovaSeq 6000 S4 in dual-index mode with eight nt indices.

### *C. parvum* RT-PCR validation of polycistrons

A total of 1 ng input *C. parvum* total RNA was treated with DNase I (Invitrogen) following the manufacturer’s protocol. Reverse transcription of RNA was carried out using SuperScript IV (Invitrogen) following the manufacturer’s protocol, using random hexamer and poly-dT primer at a 1:1 ratio. A no DNase I treated RNA tube was used as one negative control; a no RT enzyme was used as the other negative control. PCR primers targeting PT 5, PT 175 and PT 177 were designed with PrimerQuest (IDT) and were in-silico validated with Geneious Prime. The *hsp70* gene was used as a positive control. PCR was performed using the Platinum SuperFi II Green PCR Master Mix (Invitrogen) on a C1000 thermal cycler (Bio-Rad). PCR was performed using the default SuperFi II parameter for 30 cycles, with the per-cycle extension step length increased to 5 minutes. PCR results were run on a 1% agarose gel and imaged. RT-PCR primer sequences and amplicon information can be found in Supporting table 2.

#### Long-read RNA-seq processing and polycistron annotation

Iso-seq raw reads from the Sequel II platform were processed using the PacBio SMRT-Tools V11 following the default Iso-seq preset(30). Final reads in FASTQ format were aligned to *C. parvum* BGF T2T(11) genome assembly with minimap2(36). Genome-mapped reads were clustered with TAMA(74) using the high-sensitivity mode and transcript-boundary priority mode to generate transcript models. Transcript models that cover two or more ORFs in the published *C. parvum* annotation(11) are considered putative polycistrons. SQANTI3 V5.1(75) was used to filter transcript models with poor read support (with less than three reads) and poor splice junction (with less than three reads with fully matched splice donor and acceptor sites) support. Furthermore, all transcript models and genome-aligned reads were loaded as tracks into a local Apollo annotation platform(76) for manual curation and annotation. DRS raw reads from the ONT platform were processed with ONT’s Dorado V0.5.0 software using its high accuracy base-caller. Final reads in FASTQ format were processed as the Iso-seq reads were processed.

#### ATAC-seq processing, peak calling and annotation, motif analysis

Illumina Paired-End 150 ATAC-seq data sets were demultiplexed for the sporozoite and gDNA control. Read pairs were trimmed based on Q20 phred score and adapters were removed using Trim Galore. Raw reads were aligned to the genome using the “very sensitive” preset of Bowtie2 V2.5.1(77). Picard V3.0.0 was used to mark and remove optical duplicates. ATAC-seq insertion size estimation was performed with bamPEFragmentSize from the DeepTools V3.5.5 software suite(78). Genome-aligned reads for both the sporozoites and gDNA control were peaked called using MACS3 V3.0.1(48) using its Hidden Markov Model ATAC-seq caller preset (hmmr-atac flag). Peak calling is performed on gDNA ATAC-seq as control. Filtered sporozoite peaks were annotated with annotatePeaks from the HOMER V5.1 software suite. Annotated peaks were loaded into Apollo for manual curation. All putative regulatory ATAC-seq peaks were input for sequence motif enrichment analysis using the MEME V5.5.7 software suite with default parameters to present top three enriched motifs sorted by E-value. Enriched motifs were back assigned to individual peaks using the MEME output.

#### Gene Ontology (GO) enrichment and analysis

Gene IDs were input into CryptoDB.org, GO analysis tool to examine enrichment for Biological Processes, Cellular Components and Molecular Functions. Analyses were performed with the default p-value cut-off.

#### Spliced-leader sequence and trans-splicing machinery gene discovery

Spliced-leader sequences curated from published SLIDR and SLOPPR paper^53^ are used as query to perform nucleotide BLAST on *C. parvum* T2T BGF genome with local blastn. In addition, small RNA-seq reads are obtained from SRR16563270^16^ and feed into SLIDR and SLOPPR^53^ for *de novo* spliced-leader sequence discovery. The *T. brucei* trans-splicing leader sequence U2 and U5 binding protein sequences(79) (GenBank accessions: XP_843940.1 and XP_844903.1 were used as queries to perform translated nucleotide BLAST (tblastn) against the *C. parvum* T2T BGF genome locally.

#### Short-read RNA-seq analysis

The 3 replicates of 0 h and 24 h bulk RNA-seq are obtained from NCBI PRJNA530692(80), with their respective SRA numbers as follows. 0h: SRR8841053, SRR8841054, SRR8841055. 24 h: SRR8841056, SRR8841057, SRR8841058. Reads are trimmed with Trim Galore V0.6.10. Trimmed reads are aligned to the genome using STAR aligner V2.7.11b(81). Per base-pair resolution RNA expression are calculated using Deeptools V3.5.5(78) with TPM normalization and log_2_ transformation. Final bigwig files are visualized and analyzed in genome browser JBrowse V1.16.11(82).

#### Single-cell RNA-seq analysis

Data containing all single cell expression, clustering and annotation is downloaded from NCBI Gene Expression Omnibus GSE232438 as a single seurat object(52). All downstream analysis is carried out using Seurat V5.1.0(83).

## Data availability

All generated data are deposited in the National Institutes of Health – National Library of Medicine-National Center for Biotechnology Information databases and associated with BioProject PRJNA1088529

## Code Availability

Parameters for programs, and custom Python scripts used in this study are deposited in the GitHub with detailed description of code usage: https://github.com/jkissing/Xiao_Cparvum_Polycistron_2025

## Acknowledgements

This work was supported by National Institutes of Health awards R21AI144779 and R21AI80871. This study was supported in part by resources and technical expertise from the Georgia Advanced Computing Resource Center, a partnership between the University of Georgia’s Office of the Vice President for Research and Office of the Vice President for Information Technology.

## References

1. B. S. Leander, R. E. Clopton, P. J. Keeling, Phylogeny of gregarines (Apicomplexa) as inferred from small-subunit rDNA and beta-tubulin. Int J Syst Evol Microbiol 53, 345–354 (2003).

2. B. S. Leander, J. T. Harper, P. J. Keeling, Molecular phylogeny and surface morphology of marine aseptate gregarines (Apicomplexa): Selenidium spp. and Lecudina spp. J Parasitol 89, 1191–1205 (2003).

3. W. Checkley et al., A review of the global burden, novel diagnostics, therapeutics, and vaccine targets for Cryptosporidium. Lancet Infect Dis 15, 85–94 (2015).

4. I. H. Gilbert et al., Safe and effective treatments are needed for cryptosporidiosis, a truly neglected tropical disease. BMJ Glob Health 8 (2023).

5. Y. Feng, U. M. Ryan, L. Xiao, Genetic Diversity and Population Structure of Cryptosporidium. Trends Parasitol 34, 997–1011 (2018).

6. K. L. Kotloff et al., Burden and aetiology of diarrhoeal disease in infants and young children in developing countries (the Global Enteric Multicenter Study, GEMS): a prospective, case-control study. Lancet 382, 209–222 (2013).

7. S. Vinayak et al., Genetic modification of the diarrhoeal pathogen Cryptosporidium parvum. Nature 523, 477–480 (2015).

8. J. Tandel et al., Genetic Ablation of a Female-Specific Apetala 2 Transcription Factor Blocks Oocyst Shedding in Cryptosporidium parvum. mBio 10.1128/mbio.03261-22, e0326122 (2023).

9. A. Sateriale, M. Pawlowic, S. Vinayak, C. Brooks, B. Striepen, Genetic Manipulation of Cryptosporidium parvum with CRISPR/Cas9. Methods Mol Biol 2052, 219–228 (2020).

10. R. P. Baptista et al., Long-read assembly and comparative evidence-based reanalysis of Cryptosporidium genome sequences reveal expanded transporter repertoire and duplication of entire chromosome ends including subtelomeric regions. Genome Res 32, 203–213 (2022).

11. R. P. Baptista, R. Xiao, Y. Li, T. C. Glenn, J. C. Kissinger, New T2T assembly of Cryptosporidium parvum IOWA annotated with reference genome gene identifiers. bioRxiv 10.1101/2023.06.13.544219 (2023).

12. N. Yarlett, M. Morada, Long-term in vitro Culture of Cryptosporidium parvum. Bio Protoc 8, e2947 (2018).

13. G. Wilke et al., A Stem-Cell-Derived Platform Enables Complete Cryptosporidium Development In Vitro and Genetic Tractability. Cell Host Microbe 26, 123–134 e128 (2019).

14. L. Josse, A. J. Bones, T. Purton, M. Michaelis, A. D. Tsaousis, A Cell Culture Platform for the Cultivation of Cryptosporidium parvum. Curr Protoc Microbiol 10.1002/cpmc.80, e80 (2019).

15. Y. Li, R. P. Baptista, A. Sateriale, B. Striepen, J. C. Kissinger, Analysis of Long Non-Coding RNA in Cryptosporidium parvum Reveals Significant Stage-Specific Antisense Transcription. Front Cell Infect Microbiol 10, 608298 (2020).

16. Y. Li, R. P. Baptista, X. Mei, J. C. Kissinger, Small and intermediate size structural RNAs in the unicellular parasite Cryptosporidium parvum as revealed by sRNA-seq and comparative genomics. Microb Genom 8 (2022).

17. Y. Li, R. P. Baptista, J. C. Kissinger, Noncoding RNAs in Apicomplexan Parasites: An Update. Trends Parasitol 36, 835–849 (2020).

18. B. Striepen et al., Gene transfer in the evolution of parasite nucleotide biosynthesis. Proc Natl Acad Sci U S A 101, 3154–3159 (2004).

19. M. C. Pawlowic et al., Genetic ablation of purine salvage in Cryptosporidium parvum reveals nucleotide uptake from the host cell. Proc Natl Acad Sci U S A 116, 21160–21165 (2019).

20. T. Blumenthal, Operons in eukaryotes. Brief Funct Genomic Proteomic 3, 199–211 (2004).

21. A. V. Jager, J. G. De Gaudenzi, A. Cassola, I. D’Orso, A. C. Frasch, mRNA maturation by two-step transsplicing/polyadenylation processing in trypanosomes. Proc Natl Acad Sci U S A 104, 2035–2042 (2007).

22. J. A. Denker, D. M. Zuckerman, P. A. Maroney, T. W. Nilsen, New components of the spliced leader RNP required for nematode trans-splicing. Nature 417, 667–670 (2002).

23. K. B. Lidie, F. M. van Dolah, Spliced leader RNA-mediated trans-splicing in a dinoflagellate, Karenia brevis. J Eukaryot Microbiol 54, 427–435 (2007).

24. E. L. Lasda, T. Blumenthal, Trans-splicing. Wiley Interdiscip Rev RNA 2, 417–434 (2011).

25. N. Lapalu et al., Improved gene annotation of the fungal wheat pathogen Zymoseptoria tritici based on combined Iso-Seq and RNA-Seq evidence. 10.1101/2023.04.26.537486 (2023).

26. S. P. Gordon et al., Widespread Polycistronic Transcripts in Fungi Revealed by Single-Molecule mRNA Sequencing. PLoS One 10, e0132628 (2015).

27. J. Slone, J. Daniels, H. Amrein, Sugar receptors in Drosophila. Curr Biol 17, 1809–1816 (2007).

28. S. D. Gallaher et al., Widespread polycistronic gene expression in green algae. Proc Natl Acad Sci U S A 118 (2021).

29. C. H. Cho et al., Genome-wide signatures of adaptation to extreme environments in red algae. Nat Commun 14, 10 (2023).

30. M. L. Gonzalez-Garay, “Introduction to Isoform Sequencing Using Pacific Biosciences Technology (Iso-Seq)” in Transcriptomics and Gene Regulation. (2016), 10.1007/978-94-017-7450-5_6 chap. Chapter 6, pp. 141–160.

31. D. R. Garalde et al., Highly parallel direct RNA sequencing on an array of nanopores. Nat Methods 15, 201–206 (2018).

32. G. Garces-Sanchez, P. A. Wilderer, J. C. Munch, H. Horn, M. Lebuhn, Evaluation of two methods for quantification of hsp70 mRNA from the waterborne pathogen Cryptosporidium parvum by reverse transcription real-time PCR in environmental samples. Water Res 43, 2669–2678 (2009).

33. C. Swale et al., Altiratinib blocks Toxoplasma gondii and Plasmodium falciparum development by selectively targeting a spliceosome kinase. Sci Transl Med 14, eabn3231 (2022).

34. D. Puiu, S. Enomoto, G. A. Buck, M. S. Abrahamsen, J. C. Kissinger, CryptoDB: the Cryptosporidium genome resource. Nucleic Acids Res 32, D329–331 (2004).

35. M. Heiges et al., CryptoDB: a Cryptosporidium bioinformatics resource update. Nucleic Acids Res 34, D419–422 (2006).

36. H. Li, Minimap2: pairwise alignment for nucleotide sequences. Bioinformatics 34, 3094–3100 (2018).

37. C. Camacho et al., BLAST+: architecture and applications. BMC Bioinformatics 10, 421 (2009).

38. M. Kozak, Effects of intercistronic length on the efficiency of reinitiation by eucaryotic ribosomes. Mol Cell Biol 7, 3438–3445 (1987).

39. M. Kozak, Regulation of translation via mRNA structure in prokaryotes and eukaryotes. Gene 361, 13–37 (2005).

40. G. Hernandez, V. G. Osnaya, X. Perez-Martinez, Conservation and Variability of the AUG Initiation Codon Context in Eukaryotes. Trends Biochem Sci 44, 1009–1021 (2019).

41. M. Kumar, V. Srinivas, S. Patankar, Upstream AUGs and upstream ORFs can regulate the downstream ORF in Plasmodium falciparum. Malar J 14, 512 (2015).

42. J. Li, Q. Liang, W. Song, M. A. Marchisio, Nucleotides upstream of the Kozak sequence strongly influence gene expression in the yeast S. cerevisiae. J Biol Eng 11, 25 (2017).

43. E. Beaudoing, S. Freier, J. R. Wyatt, J. M. Claverie, D. Gautheret, Patterns of variant polyadenylation signal usage in human genes. Genome Res 10, 1001–1010 (2000).

44. J. D. Ospina-Villa et al., mRNA Polyadenylation Machineries in Intestinal Protozoan Parasites. J Eukaryot Microbiol 67, 306–320 (2020).

45. R. E. Workman et al., Nanopore native RNA sequencing of a human poly(A) transcriptome. Nat Methods 16, 1297–1305 (2019).

46. N. J. Loman, J. Quick, J. T. Simpson, A complete bacterial genome assembled de novo using only nanopore sequencing data. Nat Methods 12, 733–735 (2015).

47. J. D. Buenrostro, B. Wu, H. Y. Chang, W. J. Greenleaf, ATAC-seq: A Method for Assaying Chromatin Accessibility Genome-Wide. Curr Protoc Mol Biol 109, 21 29 21–21 29 29 (2015).

48. Y. Zhang et al., Model-based analysis of ChIP-Seq (MACS). Genome Biol 9, R137 (2008).

49. S. Heinz et al., Simple combinations of lineage-determining transcription factors prime cis-regulatory elements required for macrophage and B cell identities. Mol Cell 38, 576–589 (2010).

50. T. L. Bailey et al., MEME SUITE: tools for motif discovery and searching. Nucleic Acids Res 37, W202–208 (2009).

51. S. Gupta, J. A. Stamatoyannopoulos, T. L. Bailey, W. S. Noble, Quantifying similarity between motifs. Genome Biol 8, R24 (2007).

52. K. A. Walzer et al., Transcriptional control of the Cryptosporidium life cycle. Nature 630, 174–180 (2024).

53. M. A. Wenzel, B. Muller, J. Pettitt, SLIDR and SLOPPR: flexible identification of spliced leader trans-splicing and prediction of eukaryotic operons from RNA-Seq data. BMC Bioinformatics 22, 140 (2021).

54. H. Zhang et al., Spliced leader RNA trans-splicing in dinoflagellates. Proc Natl Acad Sci U S A 104, 4618–4623 (2007).

55. T. T. Mkandawire, A. Sateriale, The Long and Short of Next Generation Sequencing for Cryptosporidium Research. Front Cell Infect Microbiol 12, 871860 (2022).

56. L. M. Yeoh, V. V. Lee, G. I. McFadden, S. A. Ralph, Alternative Splicing in Apicomplexan Parasites. MBio 10 (2019).

57. C. Kaur, M. Kumar, S. Patankar, Messenger RNAs with large numbers of upstream open reading frames are translated via leaky scanning and reinitiation in the asexual stages of Plasmodium falciparum. Parasitology 147, 1100–1113 (2020).

58. C. Kaur, S. Patankar, The role of upstream open reading frames in translation regulation in the apicomplexan parasites Plasmodium falciparum and Toxoplasma gondii. Parasitology 148, 1277–1287 (2021).

59. M. A. Hassan, J. J. Vasquez, C. Guo-Liang, M. Meissner, T. Nicolai Siegel, Comparative ribosome profiling uncovers a dominant role for translational control in Toxoplasma gondii. BMC Genomics 18, 961 (2017).

60. M. Sun et al., Bicistronic design as recombinant expression enhancer: characteristics, applications, and structural optimization. Appl Microbiol Biotechnol 105, 7709–7720 (2021).

61. Y. Hirose, J. L. Manley, RNA polymerase II is an essential mRNA polyadenylation factor. Nature 395, 93–96 (1998).

62. S. A. Lima et al., Short poly(A) tails are a conserved feature of highly expressed genes. Nat Struct Mol Biol 24, 1057–1063 (2017).

63. M. Sawant et al., Putative SET-domain methyltransferases in Cryptosporidium parvum and histone methylation during infection. Virulence 13, 1632–1650 (2022).

64. T. Hollin, K. G. Le Roch, From Genes to Transcripts, a Tightly Regulated Journey in Plasmodium. Front Cell Infect Microbiol 10, 618454 (2020).

65. J. Oberstaller, Y. Pumpalova, A. Schieler, M. Llinás, J. C. Kissinger, The Cryptosporidium parvum ApiAP2 gene family: insights into the evolution of apicomplexan AP2 regulatory systems. Nucleic Acids Res 10.1093/nar/gku500 (2014).

66. J. Oberstaller, S. J. Joseph, J. C. Kissinger, Genome-wide upstream motif analysis of Cryptosporidium parvum genes clustered by expression profile. Bmc Genomics 14 (2013).

67. J. R. Nevins et al., E2F transcription factor is a target for the RB protein and the cyclin A protein. Cold Spring Harb Symp Quant Biol 56, 157–162 (1991).

68. K. Helin, Regulation of cell proliferation by the E2F transcription factors. Curr Opin Genet Dev 8, 28–35 (1998).

69. P. J. Iaquinta, J. A. Lees, Life and death decisions by the E2F transcription factors. Curr Opin Cell Biol 19, 649–657 (2007).

70. Y. Sylvestre et al., An E2F/miR-20a autoregulatory feedback loop. J Biol Chem 282, 2135–2143 (2007).

71. R. Volden et al., Improving nanopore read accuracy with the R2C2 method enables the sequencing of highly multiplexed full-length single-cell cDNA. Proc Natl Acad Sci U S A 115, 9726–9731 (2018).

72. F. Ibrahim, J. Oppelt, M. Maragkakis, Z. Mourelatos, TERA-Seq: true end-to-end sequencing of native RNA molecules for transcriptome characterization. Nucleic Acids Res 49, e115 (2021).

73. T. Chari, L. Pachter, The specious art of single-cell genomics. PLoS Comput Biol 19, e1011288 (2023).

74. R. I. Kuo et al., Illuminating the dark side of the human transcriptome with long read transcript sequencing. BMC Genomics 21, 751 (2020).

75. M. Tardaguila et al., SQANTI: extensive characterization of long-read transcript sequences for quality control in full-length transcriptome identification and quantification. Genome Res 10.1101/gr.222976.117 (2018).

76. S. E. Lewis et al., Apollo: a sequence annotation editor. Genome Biol 3, RESEARCH0082 (2002).

77. B. Langmead, S. L. Salzberg, Fast gapped-read alignment with Bowtie 2. Nat Methods 9, 357–359 (2012).

78. F. Ramirez, F. Dundar, S. Diehl, B. A. Gruning, T. Manke, deepTools: a flexible platform for exploring deep-sequencing data. Nucleic Acids Res 42, W187–191 (2014).

79. X. H. Liang, A. Haritan, S. Uliel, S. Michaeli, trans and cis splicing in trypanosomatids: mechanism, factors, and regulation. Eukaryot Cell 2, 830–840 (2003).

80. J. Tandel et al., Life cycle progression and sexual development of the apicomplexan parasite Cryptosporidium parvum. Nat Microbiol 4, 2226–2236 (2019).

81. A. Dobin et al., STAR: ultrafast universal RNA-seq aligner. Bioinformatics 29, 15–21 (2013).

82. R. Buels et al., JBrowse: a dynamic web platform for genome visualization and analysis. Genome Biol 17, 66 (2016).

83. Y. Hao et al., Dictionary learning for integrative, multimodal and scalable single-cell analysis. Nat Biotechnol 42, 293–304 (2024).

